# An integrated approach to evaluate the functional effects of disease-associated NMDA receptor variants

**DOI:** 10.1101/2022.06.02.494569

**Authors:** Gabrielle Moody, Angela Musco, Joseph Bennett, Lonnie P. Wollmuth

## Abstract

The NMDA receptor (NMDAR) is a ubiquitously expressed glutamate-gated ion channel that plays key roles in brain development and function. Not surprisingly, a variety of disease-associated variants have been identified in genes encoding NMDAR subunits. A critical first step to assess whether these variants contribute to their associated disorder is to characterize their effect on receptor function. However, the complexity of NMDAR function makes this challenging, with most variants typically altering multiple functional properties. At synapses, NMDARs encode presynaptic activity to carry a charge transfer that alters membrane excitability and a Ca^2+^ influx that has both short- and long-term signaling actions. Here, we characterized epilepsy-associated variants in GluN1 and GluN2A subunits with various phenotypic severity. To capture the dynamics of NMDAR encoding, we applied 10 glutamate pulses at 10 Hz to derive a charge integral. This encoding assay is advantageous since it incorporates multiple gating parameters – activation, deactivation, and desensitization – into a single value. We then integrated this encoding with Mg^2+^ block and Ca^2+^ influx using fractional Ca^2+^ currents to generate indices of charge transfer and Ca^2+^ transfer over wide voltage ranges. This approach yields consolidated parameters that can be used as a reference to normalize allosteric modulation and has the potential to speed up future bench to bedside methods of investigating variants to determine patient treatment.

## INTRODUCTION

Missense and nonsense mutations in diverse families of ion channels are often associated with clinical phenotypes (Marini *et al*., 2011; Noebels, 2017; Catterall, 2018; Deng & Klyachko, 2021). Characterizing how these mutations affect ion channel function is a critical step in defining their pathophysiology. Outcomes can be classified as loss-of-function (LoF) where the overall effectiveness of the protein is reduced or gain-of-function (GoF) where it is enhanced, with the specific designation providing a potential framework for treatment (Hu *et al*., 2016; Noebels, 2017; Tang *et al*., 2020).

Glutamate is the major excitatory neurotransmitter in the vertebrate nervous system. For fast signaling, glutamate is converted into a biological signal by various ionotropic glutamate receptors (iGluRs), including AMPA (AMPAR), kainate (KAR), and NMDA (NMDAR) receptor subtypes (Hansen *et al*., 2021). Recently, a variety of disease-associated missense and nonsense mutations have been identified in genes encoding iGluR subunits (Burnashev & Szepetowski, 2015; Yuan et al., 2015; Geisheker et al., 2017; Guzman et al., 2017; XiangWei et al., 2018; Xu & Luo, 2018; Garcia-Recio et al., 2021; Stolz et al., 2021). These variants are associated with numerous neurodevelopmental, neurological, and psychiatric disorders, highlighting the diverse role of iGluRs in brain development and function.

The vast majority of disease-associated iGluR variants have been identified in genes encoding NMDAR subunits. Numerous studies have characterized the effects of these variants on receptor function, though the majority remain uncharacterized (Perszyk et al., 2020; Amin et al., 2021; Garcia-Recio et al., 2021). These characterizations have provided guidelines for treatment (Pierson et al., 2014; Yuan et al., 2014; Soto et al., 2019; Xu et al., 2021). However, NMDARs have complex signaling mechanisms that are regulated by diverse biophysical properties (Paoletti et al., 2013; Hansen *et al*., 2017), and defining NMDAR variants as LoF or GoF often is challenging since they can alter these properties in opposing ways. One powerful approach to address this challenge is to integrate these diverse biophysical properties into consolidated parameters that encompass multiple biophysical properties (Swanger *et al*., 2016; Li *et al*., 2019). Here, we develop an approach to characterize disease-associated variants *in vitro* that is efficient, requiring measurement of only three parameters (charge integral with multiple glutamate applications, Mg^2+^ block, and Ca^2+^ permeation using fractional Ca^2+^ currents), and that yields consolidated descriptions of the effects of these variants on diverse biophysical properties over a wide voltage range. This approach captures aspects of the physiology of NMDARs at synapses as well as at extrasynaptic sites, potentially providing a more refined guidance for treatment options.

NMDARs are obligate heterotetramers composed of two GluN1 subunits, encoded by *GRIN1*, and typically some combination of GluN2(A-D) subunits, encoded by *GRIN2(A-D*). GluN2A is a hotspot for epilepsy, with more that 50% of variants in GluN2A having some form of epilepsy (Xu & Luo, 2018). In the present study, we characterized a set of NMDAR variants that display diverse epilepsy symptoms and severity as well as other co-morbidities (Table 1). These variants are distributed throughout the protein (Figure 1), though we included many from around the transmembrane region since these often have multiple effects on receptor function (Table 2). We chose variants with both LoF and GoF effects on the receptor, though there is often little or no data on their effects on Ca^2+^ influx, a key functional feature of NMDARs.

**Figure 1.**
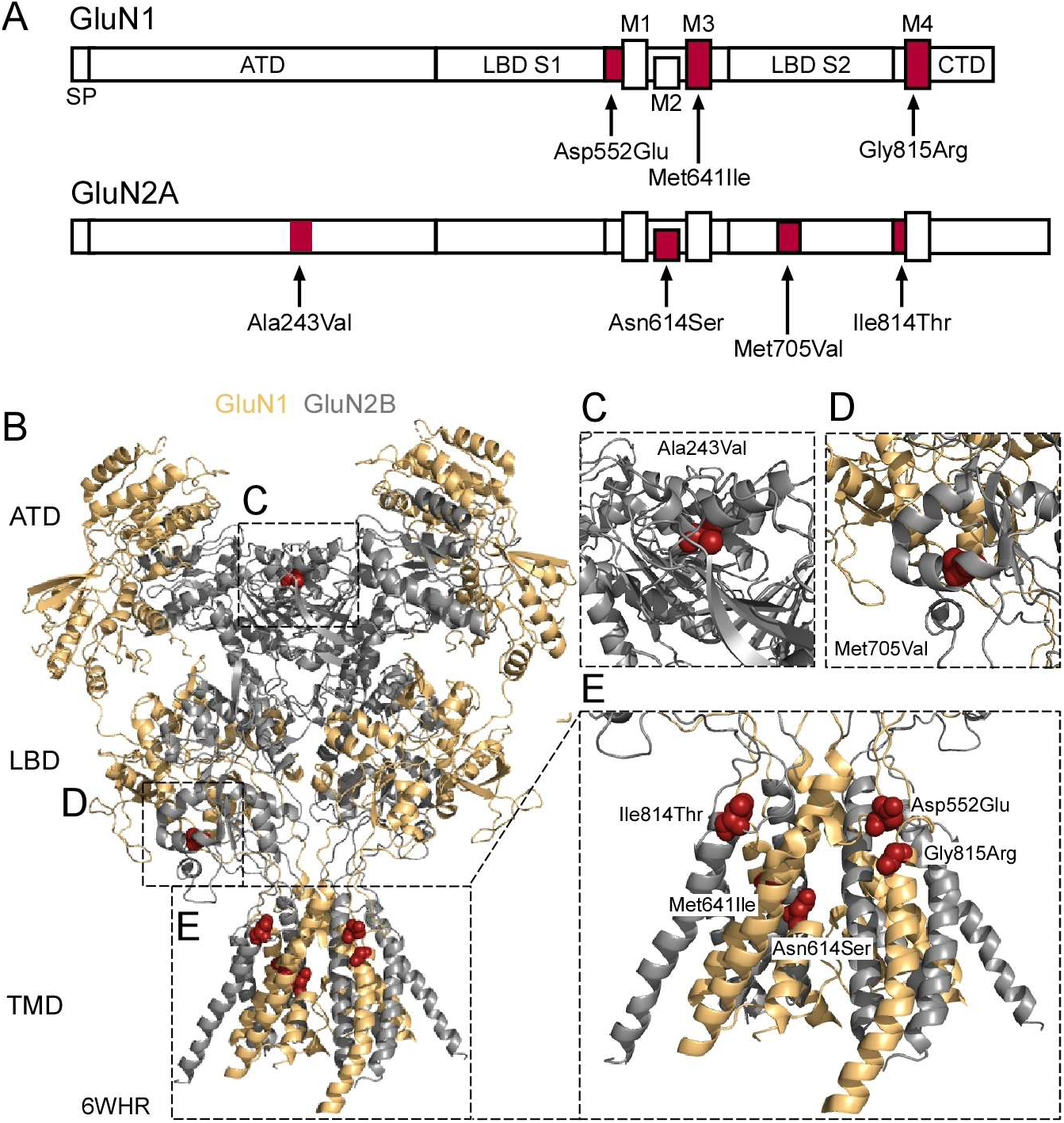
Topology of the NMDAR and distribution of tested variants. **(A)** Linear representation of GluN1 (GRIN1) and GluN2A (GRIN2A) showing distribution of tested disease-associated variants (red) (Tables 1 & 2). Variants are distributed throughout the proteins, including in the GluN1 pre-M1 (Asp552Glu, D552E), the M3 (Met641Ile, M641I) and M4 (Gly815Arg, G815R) segments, and in the GluN2A ATD (Ala243Val, A243V), M2 loop (Asn614Ser, N614S), LBD S2 (Met705Val, M705V), and pre-M4 (Ile814Thr, I814T). Abbreviations: SP, signal peptide; ATD, amino-terminal domain; LBD, ligand-binding domain lobes S1 or S2; TMD, transmembrane domain encompassing transmembrane segments M1, M3, & M4 and a M2 pore loop; CTD, C-terminal domain. **(B)** NMDARs (N1 gold, N2A gray, 6WHR (Chou et al., 2020)) have a layered topology with the ATD and LBD positioned in the synaptic cleft, the TMD or ion channel embedded in the membrane, and the CTD (not shown) located intracellularly. Variants A243V (**C)** and M705V **(D)** are in the ATD and LBD S2, respectively. Remaining variants are clustered within or around the TMD **(E)**.

**Table 1.** Selected *de novo* and inherited epilepsy-associated variants and associated clinical phenotypes.

| **GluN1** |  |  |  |  |
| --- | --- | --- | --- | --- |
| **Protein Mutation** | **# of patients** | **Age at seizure onset** | **Clinical Phenotype\*** | **Phenotype References\*\*** |
| Asp552Glu (D552E) | 3 | Birth-10 yrs | Epilepsy, severe ID/DD, ASD-like, microcephaly, hypotonia, dyskinesia, cerebral atrophy | (Ohba *et al.*, 2015; Lemke *et al.*, 2016) |
| Met641Ile (M641I) | 1 | 2 mo | Epilepsy, severe ID/DD, dyskinesia, cerebral atrophy, sleep disorder | (Ohba *et al.*, 2015; Lemke *et al.*, 2016) |
| Gly815Arg (G815R) | 3 | 2-12 mo | Epilepsy, severe ID/DD, dyskinesia, hypotonia, cerebral atrophy | (Ohba *et al.*, 2015; Lemke *et al.*, 2016) |
| **GluN2A** |  |  |  |  |
| **Protein Mutation** | **# of patients** | **Age at seizure onset** | **Clinical Phenotype\*** | **Phenotype References\*\*** |
| Ala243Val (A243V) | 1 | 5.5 yrs | BECTS/RE, LD | (Lemke *et al.*, 2013) |
| Asn614Ser (N614S) | 1 | unknown | Focal epilepsy | (von Stulpnagel *et al.*, 2017) |
| Met705Val (M705V) | 1 | 8 yrs | BECTS/RE, mild LD, speech delay | (Lemke *et al.*, 2013) |
| Ile814Thr (I814T) | 1 | 5 yrs | BECTS/RE | (Lemke *et al.*, 2013) |

Human GluN1 (GluN1; NCBI Protein database accession no. Q05586) and human GluN2A (GluN2A; Q12879) subunits. Protein number includes signal peptide (GluN1, 18 residues; GluN2A, 19 residues).
\*ID= intellectual disability, DD = developmental delay, ASD = Autism spectrum disorder, EE = epileptic encephalopathy, LD = learning difficulties/speech delays, BECTS = benign epilepsy with centrottemporal spikes/RE = Rolandic epilepsy
\*\*Earliest publication(s) only.

**Table 2.**
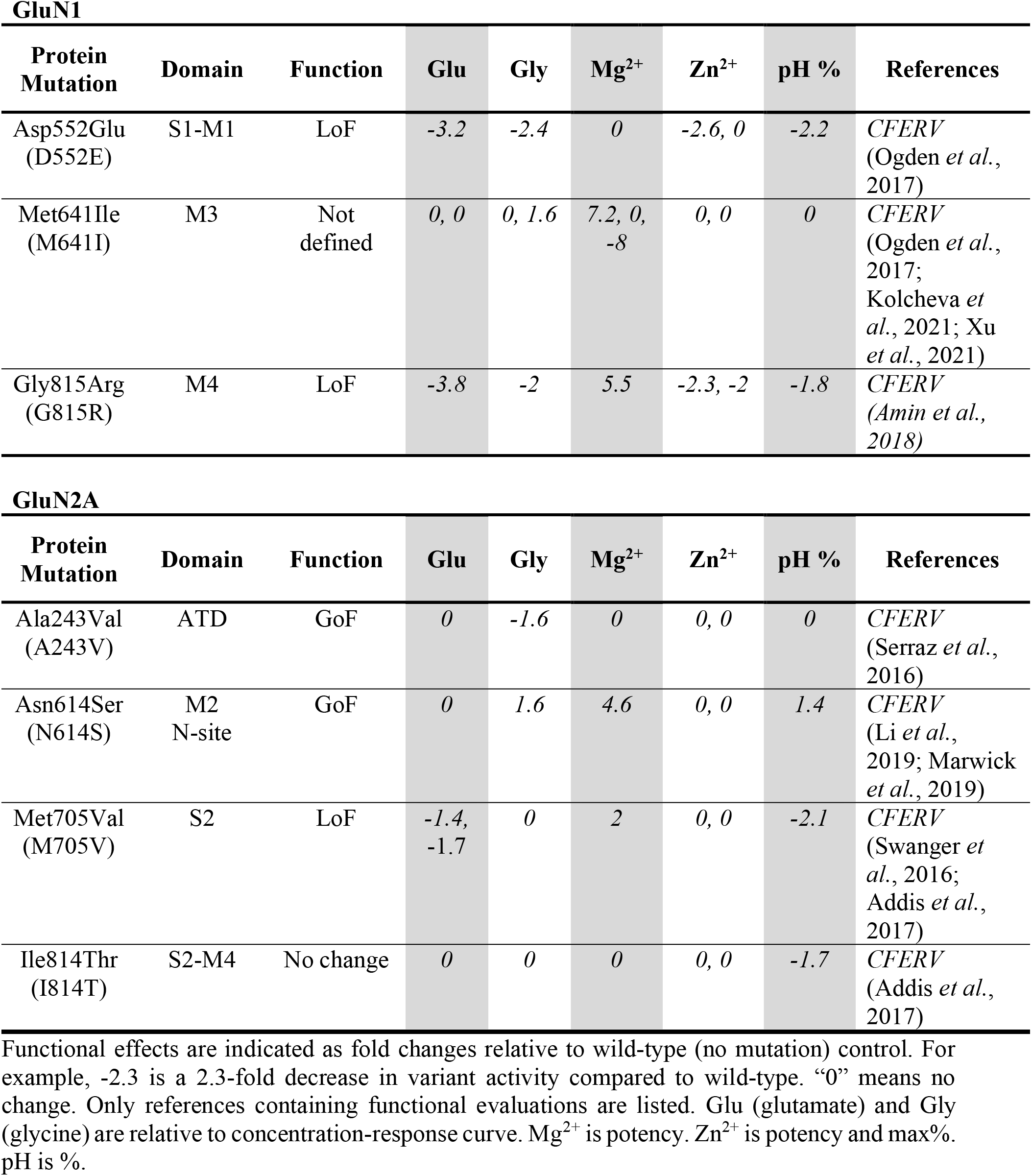
Distribution and published functional effects of NMDAR variants.

| **GluN1** |  |  |  |  |  |  |  |  |
| --- | --- | --- | --- | --- | --- | --- | --- | --- |
| **Protein Mutation** | **Domain** | **Function** | **Glu** | **Gly** | **Mg<sup>2+</sup>** | **Zn<sup>2+</sup>** | **pH %** | **References** |
| Asp552Glu (D552E) | S1-M1 | LoF | -3.2 | -2.4 | 0 | -2.6, 0 | -2.2 | *CFERV* (Ogden *et al.*, 2017) |
| Met641Ile (M641I) | M3 | Not defined | 0, 0 | 0, 1.6 | 7.2, 0, -8 | 0, 0 | 0 | *CFERV* (Ogden *et al.*, 2017; Kolcheva *et al.*, 2021; Xu *et al.*, 2021) |
| Gly815Arg (G815R) | M4 | LoF | -3.8 | -2 | 5.5 | -2.3, -2 | -1.8 | *CFERV* (Amin *et al.*, 2018) |
| **GluN2A** |  |  |  |  |  |  |  |  |
| **Protein Mutation** | **Domain** | **Function** | **Glu** | **Gly** | **Mg<sup>2+</sup>** | **Zn<sup>2+</sup>** | **pH %** | **References** |
| Ala243Val (A243V) | ATD | GoF | 0 | -1.6 | 0 | 0, 0 | 0 | *CFERV* (Serraz *et al.*, 2016) |
| Asn614Ser (N614S) | M2 N-site | GoF | 0 | 1.6 | 4.6 | 0, 0 | 1.4 | *CFERV* (Li *et al.*, 2019; Marwick *et al.*, 2019) |
| Met705Val (M705V) | S2 | LoF | -1.4, -1.7 | 0 | 2 | 0, 0 | -2.1 | *CFERV* (Swanger *et al.*, 2016; Addis *et al.*, 2017) |
| Ile814Thr (I814T) | S2-M4 | No change | 0 | 0 | 0 | 0, 0 | -1.7 | *CFERV* (Addis *et al.*, 2017) |

Functional effects are indicated as fold changes relative to wild-type (no mutation) control. For example, -2.3 is a 2.3-fold decrease in variant activity compared to wild-type. “0” means no change. Only references containing functional evaluations are listed. Glu (glutamate) and Gly (glycine) are relative to concentration-response curve. Mg<sup>2+</sup> is potency. Zn<sup>2+</sup> is potency and max%. pH is %.

A hallmark of synaptic NMDARs is their ability to integrate multiple synaptic release events and transduce them into patterns of neuronal activity (Hansen *et al*., 2021). This NMDAR frequency response or ‘encoding’ impacts two general signaling mechanisms of NMDARs: a charge transfer, mediated by Na^+^ & Ca^2+^ which impacts membrane excitability (Schiller *et al*., 2000; Palmer et al., 2014; Stuart & Spruston, 2015), and a Ca^2+^ influx that can affect local events such as synapse structure and strength of signaling (Malenka & Nicoll, 1999; Citri & Malenka, 2008; Herring & Nicoll, 2016), distal events such as gene expression (Bading, 2013), and when unregulated cell viability (Choi, 2020).

As an initial step to contrast NMDAR variants with wild-type, we evaluated gating by deriving a charge transfer from multiple synaptic-like applications of glutamate. This approach is advantageous since it encompasses multiple biophysical properties including receptor activation, deactivation, and desensitization within a single charge transfer term and is more representative of how NMDARs function at synapses than single glutamate applications. Subsequently, we corrected the voltage dependence of this charge transfer for variant-induced alterations in Mg^2+^ block. Lastly, we measured Ca^2+^ influx under physiological conditions using fractional Ca^2+^ currents (Neher, 1995; Jatzke et al., 2002). From these quantifications, we can derive a charge and Ca^2+^ transfer terms over wide voltage ranges and use appropriate allosteric modulator to normalize these parameters to that for wild-type.

## MATERIALS AND METHODS

### Molecular biology and expression

All manipulations were performed in human GluN1 (hGluN1; NCBI Protein database accession no. Q05586) and human GluN2A (hGluN2A; Q12879) subunits. In all cases, numbering included the signal polypeptide (hGluN1, 18 residues; hGluN2A, 19 residues). Constructs were generated using site-directed mutagenesis via QuickChange (Agilent) with XL1-Blue super-competent cells (primer sequences and sequencing are available upon request). Molecular structures were visualized in PyMOL (The PyMOL Molecular Graphics System, Version 1.8.4.0 Schrödinger, LLC).

Human embryonic kidney 293 (HEK293) cells were grown in Dulbecco’s modified Eagle’s medium (DMEM) with 10% fetal bovine serum (FBS) for 24 h preceding transfection (Alsaloum *et al*., 2016; Amin *et al*., 2017). Non-tagged cDNA constructs in a PCI-neo vector and a pEGFP-Cl vector (200 ng/μL) were transfected into HEK293 cells at a ratio of 3:3:1 (hGluN1/hGluN2A/EGFP) using X-tremegene HP (Roche) at a 1:100 ratio with optiMEM. Four hours after transfection, the optiMEM transfection media was replaced with 5% FBS DMEM culture media containing APV (100 μM) and Mg^2+^ (1 mM). Experiments were performed 18-36 hours after transfection.

### Whole-cell current recordings

All macroscopic currents were recorded in the whole-cell mode from HEK293 cells at room temperature (20-23°C) using an EPC-10 amplifier and Patchmaster software (HEKA Elektronic, Lambrecht, Germany), digitized at 10 kHz and low-pass Bessel filtered at 2.9 kHz. Patch microelectrodes were filled with a KCl-based intracellular solution (in mM): 140 KCl, 2 NaCl, 4 Mg^2+^-ATP, 0.3 Na^+^-ATP, 1 BAPTA, 10 HEPES, pH 7.2 (KOH), 300 mOsm (sucrose) and had resistances of 4-6 MΩ. Our standard external solution contained (in mM): 150 NaCl, 2.5 KCl, 10 HEPES, pH 7.2 (NaOH). On the day of recording, 1 mM CaCl2 and 0.1 mM glycine were added to external solutions unless otherwise noted. 1 mM glutamate was added to the agonist solution. Both glycine and glutamate were at saturating concentrations. A piezo-driven double barrel application system was used to apply external solutions with one barrel containing our external solution plus 0.1 mM glycine and the other barrel containing the same solution plus 1 mM glutamate (Yelshansky *et al*., 2004). The 10-90% rise time of the application pipet was determined at the start of each recording cycle using 10% external solution. The open tip response was 400-600 μs.

### Data collection and analysis

Wild-type recordings were made on the same transfection cycle as the variants. Recordings with changes in resistance greater than 500 MΩ were not included in analysis. All analysis was performed using custom written programs in IgorPro (WaveMetrics Inc., Lake Oswego, OR).

#### Capacitance

We measured membrane capacitance (C_m_) in Patchmaster once the cell was lifted and placed in front of the application pipette. Because of distortions of cells arising from the perfusion system, we only included in analysis those cells that showed a capacitance from 5 pF to 20 pF. These limits were determined assuming an average diameter of a HEK293 cell is 15-20 μm and a specific membrane capacitance of 1 μF/cm^2^ (Blumlein *et al*., 2017; Bandmann *et al*., 2019).

#### Rapid glutamate pulses

To determine the frequency response or encoding of NMDARs, we rapidly applied 10 pulses of glutamate (2 ms each) at a rate of 10 Hz in the whole-cell mode. The external solution contained no Mg^2+^. For a subset of recordings, we also recorded the frequency response at −90, −60, −30, and +30 mV to determine the voltage dependence of current integrals.

For each frequency response, we measured the current integral to determine the charge transfer (pQ), the peak current amplitude for each pulse (I_peakX_, where X is the pulse number), and the decay of current amplitudes after the last pulse. We normalized this charge transfer either to C_m_ (pQ/pF), which would tell us how the variant might affect the total integral, or to I_peak1_ (pQ/pA), which would tell us how the gating properties of the receptor affected its ability to integrate multiple glutamate pulses. We also derived ΔI_peak_, which is the percent change in peak current amplitude from the 1^st^ to the last peak current peak (ΔI_peak_ = 100 * (1 - I_peak10_/I_peak1_). We fit a double exponential to the decay of currents after the last pulse to derive the weighted tau (τ_weighted_), which is the weighted sum of the individual components of a double exponential fit to current decay. We used ΔI_peak_ as an index of desensitization and decay (τ_weighted_) as an index of deactivation.

#### Extracellular Mg^2+^ block

Mg^2+^ block was recorded and analyzed using a linearization approach as done previously (Wollmuth *et al*., 1998). We used this approach since we could assay the block with a single Mg^2+^ concentration and then could model the block using derived parameters with any Mg^2+^ concentration at any membrane voltage. Briefly, a current-voltage relationship (I-V) for peak glutamate-activated currents was generated from −120 to +60 mV, initially in our standard external solution (baseline I-V), then in the same solution with added 0.1 mM Mg^2+^ (Mg^2+^ I-V), and then back to our standard external solution (washout I-V). We used a moderate Mg^2+^ concentration (0.1 mM) since it allowed a more dynamic range of the block to be quantified (Wollmuth *et al*., 1998). For both the Mg^2+^ and washout recordings, we would make multiple I-V recordings to ensure complete solution exchange.

The linearization approach is sensitive to non-Mg^2+^ induced changes in the I-V relationships. We therefore compared the overall shape of the baseline and washout I-Vs to ensure that they were generally linear and also looked at the reversal potential for each I-V relationship to ensure that these did not change dramatically or that any change was linear (the reversal potential in Mg^2+^ was the mean of the reversal potentials for the baseline and washout I-Vs). For records included in analysis, we averaged the baseline and washout I-Vs.

To quantify Mg^2+^ block, we started with the Woodhull model (Woodhull, 1973), where current amplitudes in the presence (I_Blocked_ or I_B_) and absence (no added Mg^2+^ or I_0_) of Mg^2+^ are related by:

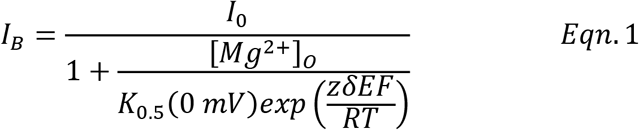

where δ is the portion of the transmembrane electric field sensed by the blocking site, K_0.5_(0 mV) is the half-maximal block at 0 mV, and z is the valence of the blocking ion. R, T, and F have their normal thermodynamic meanings and the quantity RT/F was 25.4 mV (21°C).

To determine δ and K_0.5_(0 mV), we linearized Equation 1 (DiFrancesco, 1982; Wollmuth et al., 1998):

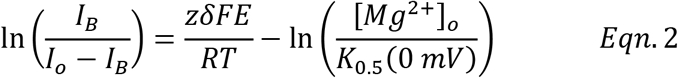

and plotted ln(I_B_/(I_0_ – I_B_)), referred to as (ln(r)), against voltage. The linear region of such a plot follows the assumptions of the Woodhull model with deviations at negative potentials most likely reflecting weak Mg^2+^ permeation (Wollmuth et al., 1998). The approximate linear region for each construct was determined using average plots of ln(r) against voltage (using a maximal ln(r) of 2.5, i.e., where the block was at least 10%) and then the same approximate voltage range was used to fit individual records.

#### Fractional Ca^2+^ currents

Fluorescence measurements were obtained as previously described (Wollmuth & Sakmann, 1998; Jatzke *et al*., 2002). Briefly, an internal solution containing 1 mM Fura-2 (K5-Fura-2, cell impermeable, Invitrogen F1200), 140 mM KCl, 10 mM HEPES, pH 7.2 was loaded into HEK293 cells via the patch pipette. The external solutions contained 140 mM NaCl, 10 mM HEPES, pH 7.2, and 1-5 mM Ca^2+^ as indicated. Cells were centered in identical positions using a camera for guidance (FLIR, SpinMaster software) and were illuminated alternatively at excitation wavelengths of 340 nm and 385 nm (10 Hz) by an optronic beam combiner (Lambda421, Sutter). Fluorescence shutter and intensity were controlled through Patchmaster. Emission fluorescence was detected using a photomultiplier and digitized in Patchmaster.

To assay Ca^2+^ influx and depending on current amplitudes, we applied a 250 to 1000 ms glutamate pulse or a pulse train of 10 pulses at 10 Hz, totaling 1000 ms, to NMDAR-expressing HEK293 cells. We minimized the duration of the glutamate application based on current amplitudes to limit Ca^2+^ influx to stay in the linear range of Fura-2. We quantified fractional Ca^2+^ currents (P_f_) using:

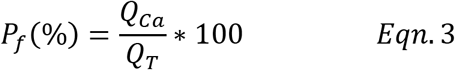

where Q_T_ is the total charge determined by the current integral of a given glutamate pulse and Q_Ca_ is the charge carried by Ca^2+^ derived from:

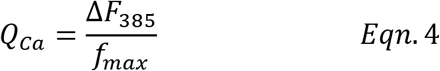

Q_Ca_ was derived from the change in F_380_ (ΔF_380_) arising from the glutamate application and f_max_, which was determined by Ca^2+^ only currents through the NMDAR at −100 mV using an external solution consisting of 10 mM Ca^2+^, 140 mM N-methyl-D-glucamine (NMDG^+^), which does not permeate the receptor, 10 mM HEPES, and our standard KCl-based internal solution (Wollmuth & Sakmann, 1998; Jatzke et al., 2002; Watanabe *et al*., 2002).

We normalized fluorescence intensity for each day based on bead units (BU), which were 4.5 mm diameter BB beads (Fluoresbrite™). The BU was determined at the end of the day as the mean fluorescence of 5–10 beads at 385 nm excitation at a set gain. All membrane potentials reported for P_f_ measurements are relative to the reversal potential. Analysis was done custom written programs in Igor Pro (WaveMetrics Inc., Lake Oswego, OR, USA).

To interelate P_f_s and P_Ca_/P_Na_, we followed the approach as described in detail previously (Schneggenburger, 1996; Wollmuth & Sakmann, 1998). Briefly, we related P_f_s to P_Ca_/P_Na_ assuming the Goldman-Hodgkin-Katz (GHK) assumptions by the relationship:

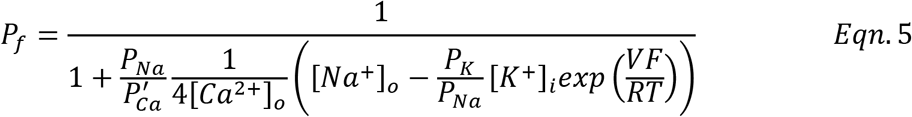

Where P_Ca_’ = P_Ca_(1 + exp(VF/RT)) and R, T, and F have their normal thermodynamic meanings. RT/F was 25.4 mV (21°C). V refers to the membrane potential relative to the reversal potential (V = V_test_ – V_rev_) where V_test_ and V_rev_ are the test and reversal potentials, respectively. V_rev_ was tested for each cell. Potentials were not corrected for junction potentials. We assumed P_Na_/P_K_ = 1.

#### Negative (NAM) and positive (PAM) allosteric modulators

NAMs or PAMs or their respective control were added to our standard external solution. The NAM Dextrorphan (DEX, Sigma), the active metabolite of Dextromethorphan, was dissolved in 1:10 DMSO and saline (140 mM NaCl) at a stock concentration of 3 mM. A final concentration of 0.3 or 3 μM was used for experiments. The control solution contained 0.01% DMSO.

The non-selective PAM GNE-9278 (Sigma) was dissolved in DMSO at a stock concentration of 30 mM. A final concentration of 30 μM was used for experiments. The control solution for GNE-9278 contained 1% DMSO.

Recordings were started in the respective DMSO control solution, where we generated a baseline glutamate pulses (10 pulses at 10 Hz). Subsequently, we made pulse recordings (2.5 s glutamate applications) before and immediately after starting wash-in (0 min) and were used to monitor drug wash-in over 2 min (15 sweeps). Sweeps 1-3 and sweeps 13-15 were used as quality control. We then generated glutamate pulses (10 pulses at 10 Hz) at 3, 4, or 5 minutes post washin. For most traces, 3 min and 5 min showed no difference. Neither drug could be washed out within a 5-10 min period.

### Statistics

All curve fitting was done using Igor Pro (WaveMetrics Inc., Lake Oswego, OR, USA). Data analysis was performed using IgorPro, Excel, and R Studio. Average values are presented as mean ± SEM. The number of replicates is indicated in the figure legend or in a table associated with the figure. Since we were specifically interested in the relationship of the variants to wild-type, we mainly carried out *Student t-tests* to determine significance., In instances where we were interested in the relationship between the variants, we used analysis of variance (ANOVA), followed by the Tukey or Dunnett’s multiple comparisons.

## RESULTS

At individual release sites, presynaptic activity is defined by action potential firing and release probability. Postsynaptic NMDARs will encode this presynaptic activity and, when open, will carry a charge that impacts electric signaling and a Ca^2+^ influx. To begin to address these dynamic aspects of NMDAR-mediated signaling and how variants might impact it, we characterized the response of wild-type and NMDAR variants to multiple glutamate pulses.

### Trains of glutamate pulses capture the dynamics of NMDAR charge transfer

Neurons fire action potentials over a wide range, from several Hz to greater than 100 Hz (Buzsaki & Mizuseki, 2014). To obtain a snapshot of NMDAR encoding, we applied 10 brief glutamate pulses (2 ms) at 10 Hz to NMDARs in the whole-cell mode (Figure 2). This frequency would approximate a presynaptic neuron firing, for example, at 10 Hz with a release probability of 1, at 20 Hz with a release probability of 0.5, or at 40 Hz with a release probability of 0.25.

**Figure 2.**
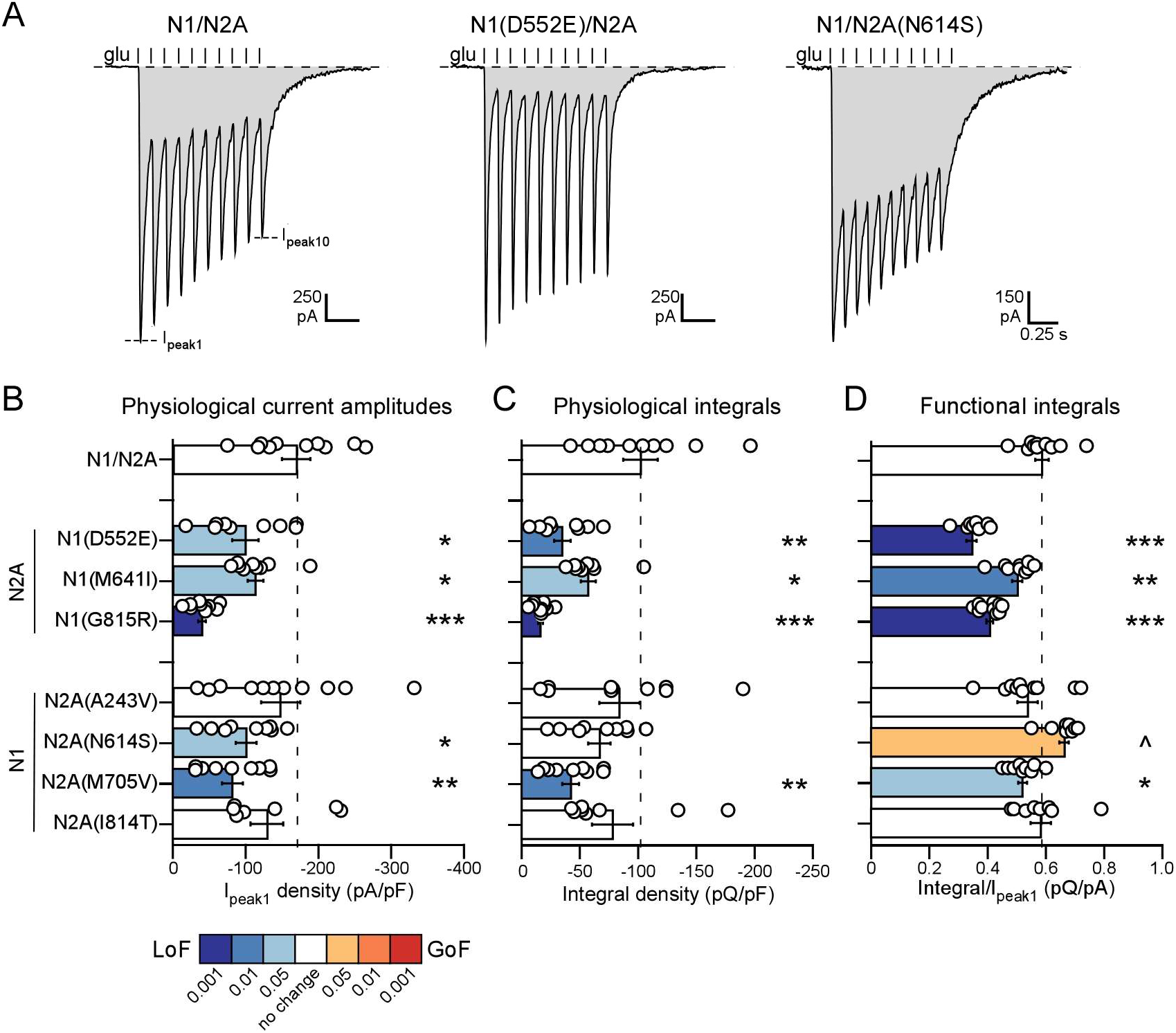
Variants alter NMDAR encoding of multiple glutamate pulses. **(A)** Whole-cell currents elicited by rapid 2 ms applications of glutamate (1 mM, vertical lines) at 10 Hz in the continuous presence of glycine (0.1 mM) as occurs at synapses. HEK293 cells were transfected with human GluN1 (N1) and human GluN2A (N2A) or with specific variants. From these recordings, we derived the current integral (shaded light gray), which varied between the constructs. Holding potential, −60 mV. **(B-D)** Bar graphs (mean ± SEM, dots indicate individual values) showing amplitude of the first glutamate application (I_peak1_) (**B**) or current integral (**C**) normalized to membrane capacitance (Integral density), which corrects for variations in cell size, or current integral normalized to I_peak1_ (**D**), which corrects for variations in the initial peak current. Values are significantly less (*p < 0.05, **p < 0.01, ***p < 0.001) or greater (^p < 0.05) than wild-type, Student t-test (Table 2). Bars are colored to match significance in terms of LoF or GoF (see Inset).

For wild-type GluN1/GluN2A, this glutamate stimulation produced a train of inward current peaks that decayed in amplitude with each successive pulse (Figure 2A, *left*). We used the integral of the current response (gray) to derive the charge transfer, which for wild-type was around −780 pQ (−780 ± 110 pQ, n = 12; mean ± SEM, n = number of whole-cell recording) (Table 3). Similar stimulations were made on the NMDAR variants (e.g., Figure 2A, *middle* & *right*).

**Table 3.**
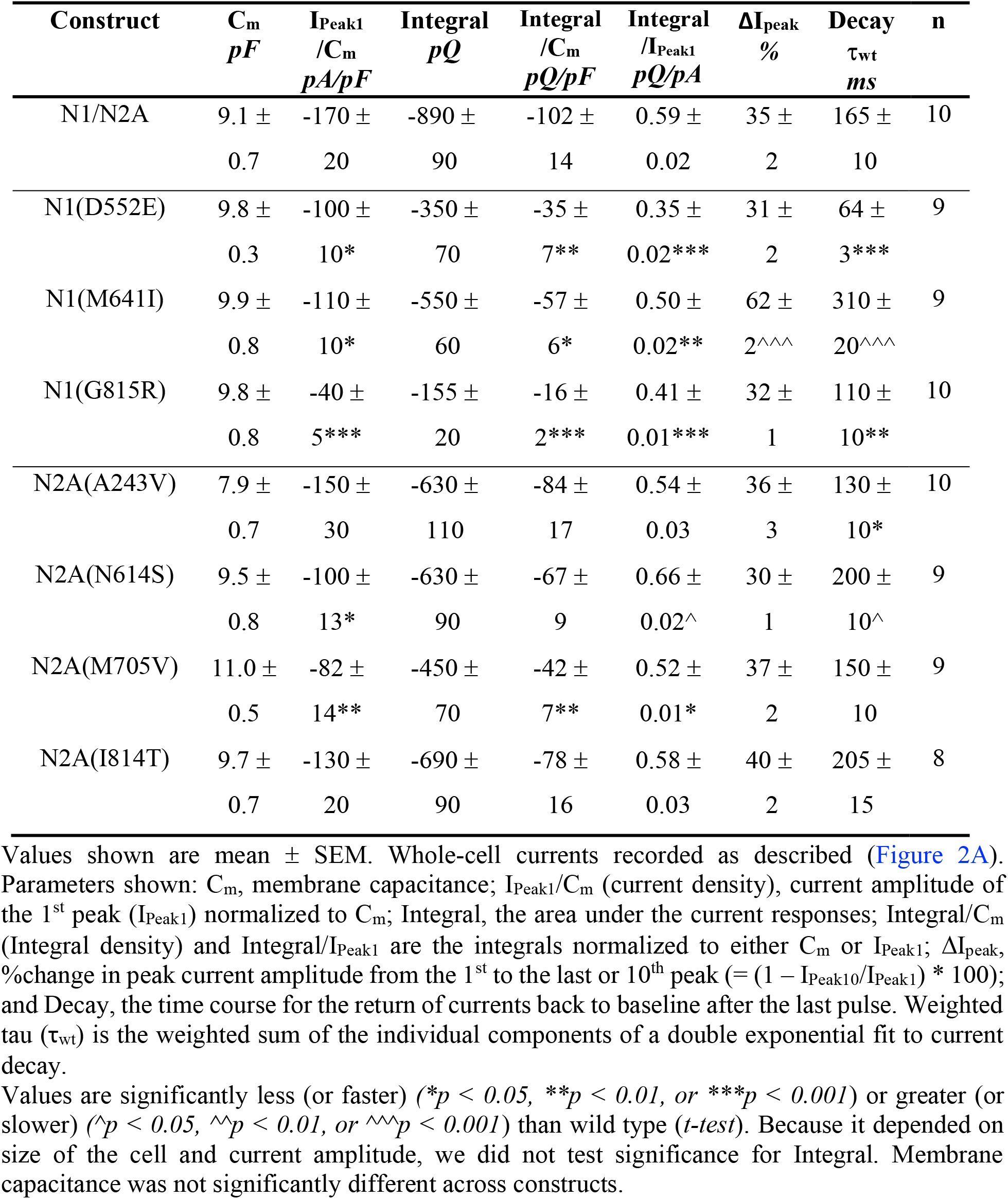
Current responses of wild-type N1/N2A and NMDAR variants to multiple glutamate applications.

| Construct | C<sub>m</sub><br>pF | I<sub>Peak1</sub><br>/C<sub>m</sub><br>pA/pF | Integral<br>pQ | Integral<br>/C<sub>m</sub><br>pQ/pF | Integral<br>/I<sub>Peak1</sub><br>pQ/pA | ΔI<sub>peak</sub><br>% | Decay<br>τ<sub>wt</sub><br>ms | n |
| --- | --- | --- | --- | --- | --- | --- | --- | --- |
| N1/N2A | 9.1 ±<br>0.7 | -170 ±<br>20 | -890 ±<br>90 | -102 ±<br>14 | 0.59 ±<br>0.02 | 35 ±<br>2 | 165 ±<br>10 | 10 |
| N1(D552E) | 9.8 ±<br>0.3 | -100 ±<br>10* | -350 ±<br>70 | -35 ±<br>7** | 0.35 ±<br>0.02*** | 31 ±<br>2 | 64 ±<br>3*** | 9 |
| N1(M641I) | 9.9 ±<br>0.8 | -110 ±<br>10* | -550 ±<br>60 | -57 ±<br>6* | 0.50 ±<br>0.02** | 62 ±<br>2^^^ | 310 ±<br>20^^^ | 9 |
| N1(G815R) | 9.8 ±<br>0.8 | -40 ±<br>5*** | -155 ±<br>20 | -16 ±<br>2*** | 0.41 ±<br>0.01*** | 32 ±<br>1 | 110 ±<br>10** | 10 |
| N2A(A243V) | 7.9 ±<br>0.7 | -150 ±<br>30 | -630 ±<br>110 | -84 ±<br>17 | 0.54 ±<br>0.03 | 36 ±<br>3 | 130 ±<br>10* | 10 |
| N2A(N614S) | 9.5 ±<br>0.8 | -100 ±<br>13* | -630 ±<br>90 | -67 ±<br>9 | 0.66 ±<br>0.02^ | 30 ±<br>1 | 200 ±<br>10^ | 9 |
| N2A(M705V) | 11.0 ±<br>0.5 | -82 ±<br>14** | -450 ±<br>70 | -42 ±<br>7** | 0.52 ±<br>0.01* | 37 ±<br>2 | 150 ±<br>10 | 9 |
| N2A(I814T) | 9.7 ±<br>0.7 | -130 ±<br>20 | -690 ±<br>90 | -78 ±<br>16 | 0.58 ±<br>0.03 | 40 ±<br>2 | 205 ±<br>15 | 8 |

Values are significantly less (or faster) (\**p* < 0.05, \*\**p* < 0.01, or \*\*\**p* < 0.001) or greater (or slower) (^*p* < 0.05, ^^*p* < 0.01, or ^^*p* < 0.001) than wild type (*t*-test). Because it depended on size of the cell and current amplitude, we did not test significance for Integral. Membrane capacitance was not significantly different across constructs.

To consider how the variants might impact NMDAR signaling, we characterized their effect on the initial peak current amplitude (I_peak1_) (Figure 2B) and current integral (Figure 2C) normalized to membrane capacitance (C_m_) (Table 3). I_peak1_/C_m_ or integral/C_m_ corrects for variations in cell size and are indices of relative membrane expression. Five of the variants, N1(D552E), N1(M641I), N1(G815R), N2A(N614S), and N2A(M705V) showed significantly reduced initial peak current amplitudes (Figure 2B). Physiological integrals showed a similar albeit not identical pattern with four variants showing statistical significance (Figure 2C). Since our focus is on charge transfer, we classified these variants as LoF in terms of the overall effectiveness of the receptor.

The current response of wild-type and the variants reflects their ability to encode the glutamate stimulations. To assay variations in this encoding ability, we normalized charge transfers to the peak amplitude of the initial glutamate stimulation (I_peak1_) (Figure 2D, Table 2). Integral/I_peak1_ normalizes the integral to the initial current amplitude and would account for variations in the ability of different construct to summate the glutamate pulses. With this normalization, wild-type had a charge transfer of 0.56 ± 0.02 pQ/pA. All GluN1 variants showed a significantly decreased charge transfer, whereas for GluN2, the charge transfer was significantly increased for N2A(N614S) and decreased for N2A(M705V).

The mechanistic basis for these variations in NMDAR encoding, as defined by Integral/I_peak1_, presumably entails variations in NMDAR gating. For example, N1(D552E) showed a more rapid deactivation time than wild-type (Decay, Table 3), and this presumably underlies the reduced charge transfer (Figure 2A, *middle*), whereas N2A(N614S) showed a significantly slower deactivation time, which presumably contributes to the increased charge transfer (Figure 2A, *right*). However, deactivation rates alone cannot account for variations in charge transfer since N1(M641I) had a significantly slower deactivation rate yet showed a decreased charge transfer. This decreased charge transfer presumably reflects an enhanced desensitization since N1(M641I) showed significantly reduced peak current amplitudes during the stimulation (ΔI_peak_, Table 3). While this interplay between deactivation and desensitization as well as other gating parameters is extremely intriguing and highly relevant to NMDAR encoding, we will not explore it further here. Still, this outcome highlights that charge transfer as an index of variants encompasses multiple gating properties of NMDARs within a single parameter.

In summary, based on charge transfer with a stimulation of 10 pulses at 10 Hz, four of the NMDAR variants produced a significant reduction in physiological charge transfer (Figure 2C). We classify these variants as LoF in terms of the effectiveness of receptor signaling.

### Voltage dependence of current integrals

NMDARs experience a wide range of voltages, which will impact current flow due to changes in driving force and the extent of Mg^2+^ block. To capture voltage-dependent features, we characterized current integrals over a wide voltage range (Figure 3) and integrated them with Mg^2+^ block (see below).

**Figure 3.**
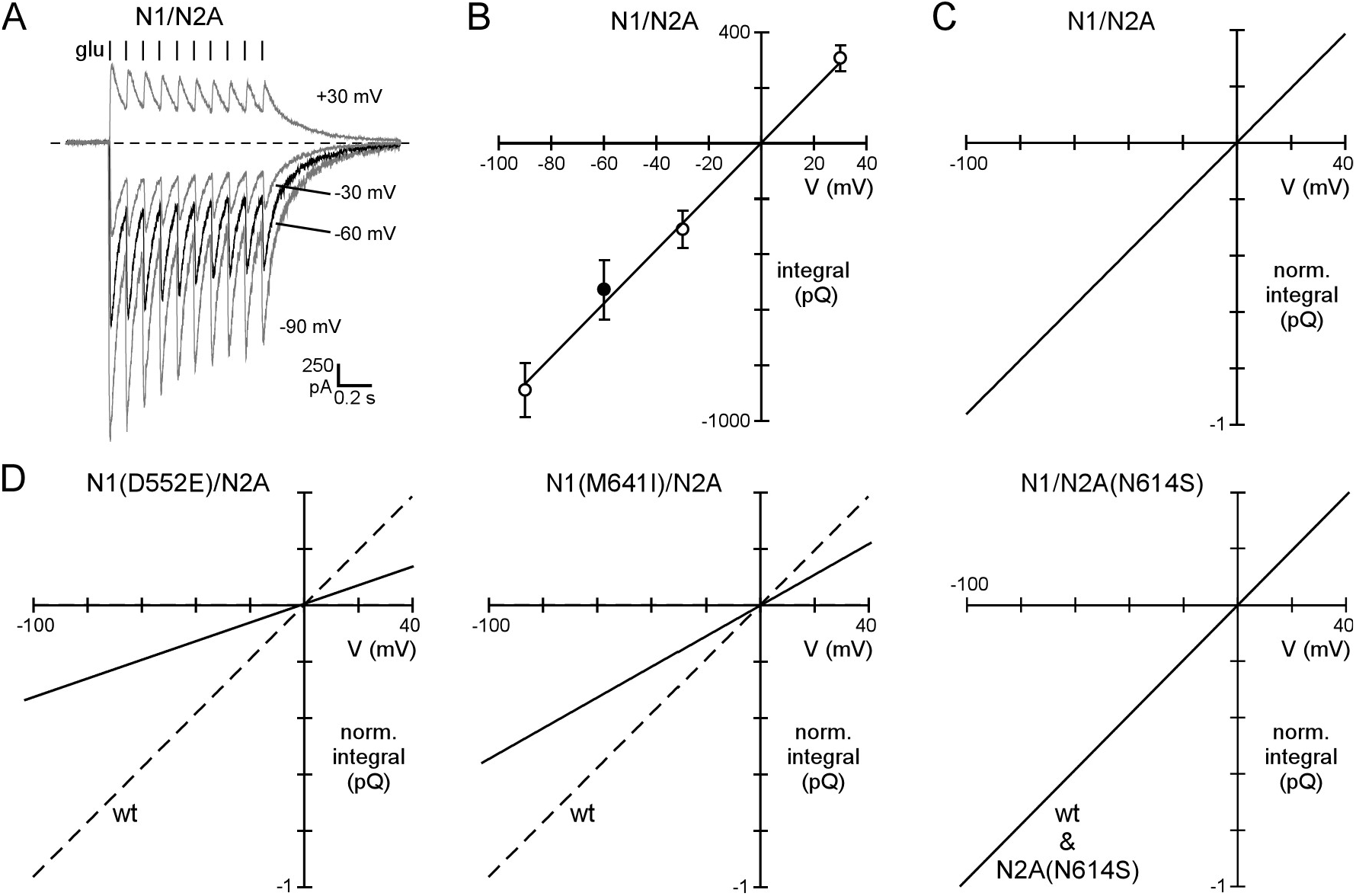
Integral-voltage (Q-V) relationships. (**A & B**) Current integrals change linearly with voltage. (**A**) Example records of current responses to 10 glutamate applications at 10 Hz for N1/N2A measured at different membrane potentials. (**B**) Integral-voltage (Q-V) relationship. Values shown are mean ± SEM (n = 5 for each; same 5 cells were recorded at each voltage). The line is a linear fit to the data with y-intercept set to 0 (slope = 9.65, R^2^ = 0.99). (**C & D**) Normalized Q-V relationships for wild-type (**C**), which is the fitted line in (**B**) normalized to the value at −100 mV, and for 3 variants (**D**). All variants showed a linear Q-V relationship (Table 4), and we used the Integral densities (Figure 2C) to normalize their Q-V relationship relative to wild-type because of the larger number of recordings. The Q-V relationship was set to wild-type (e.g., N2A(N614S)) if the Integral density was not significantly different (Table 4).

To define current integrals over a wide voltage range, we characterized them at four different potentials, −90, −60, −30, and +30 mVs for wild-type (Figure 3A) and all variants (Table 4). Like wild-type (Figure 3B), all variants showed a linear (ohmic) integral-voltage (Q-V) relationship. We normalized the Q-V relationship for wild-type (Figure 3C) and each of the variants relative to wild-type (Figures 3D) using the relative physiological integral (Figure 2C & Table 4). For each variant, we used these Q-V relationships, normalized to wild-type, as a 0 mM Mg^2+^ reference. For variants that did not show a significant change in the relative physiological integral, we reverted the value back to wild-type (e.g., N2A(N614S)), Figure 3D, *far right*).

**Table 4.** Charge-voltage curves and net charge transfer for wild-type N1/N2A and NMDARs variants.

| Construct | R<sup>2</sup> | n | Integral density ratios (-60 mV) | Net charge transfer (pQ) | -25/-70 mV current ratio | n |
| --- | --- | --- | --- | --- | --- | --- |
| N1/N2A | 0.99 | 5 | 1.00 ± 0.19 | 3.9 ± 0.5 | 4.9 ± 0.4 | 9 |
| N1(D552E)/N2A | 0.98 | 4 | 0.34 ± 0.08** | 1.5 ± 0.1*** | 3.5 ± 0.5 | 5 |
| N1(M641I)/N2A | 0.98 | 3 | 0.56 ± 0.10* | 6.2 ± 0.5^ | 1.6 ± 0.1*** | 5 |
| N1(G815R)/N2A | 0.99 | 4 | 0.16 ± 0.03*** | 1.2 ± 0.1*** | 1.3 ± 0.2*** | 7 |
| N1/N2A(A243V) | 0.99 | 3 | 0.82 ± 0.20 | 5.3 ± 0.3^ | 3.8 ± 0.4 | 5 |
|  |  |  | 1 |  |  |  |
| N1/N2A(N614S) | 0.96 | 4 | 0.65 ± 0.13 | 11.4 ± 0.7^^^ | 0.8 ± 0.1*** | 6 |
|  |  |  | 1 |  |  |  |
| N1/N2A(M705V) | 0.98 | 4 | 0.41 ± 0.09** | 1.4 ± 0.1*** | 4.4 ± 0.2 | 6 |
| N1/N2A(I814T) | 0.97 | 5 | 0.77 ± 0.19 | 3.1 ± 0.3 | 4.8 ± 0.1 | 5 |
|  |  |  | 1 |  |  |  |

### NMDAR variants alter Mg^2+^ block

One of the hallmarks of NMDARs is the regulation of their charge transfer and Ca^2+^ influx by a strong voltage-dependent block by extracellular Mg^2+^ (Mayer *et al*., 1984; Nowak *et al*., 1984). This Mg^2+^ block plays physiological roles in regulating plasticity (Citri & Malenka, 2008) and is often disrupted in NMDAR variants, especially those associated with the M2 loop (Fedele *et al*., 2018; Li *et al*., 2019; Marwick *et al*., 2019). We therefore characterized Mg^2+^ block in wild-type and the NMDAR variants (Figure 4; Table 5).

**Figure 4.**
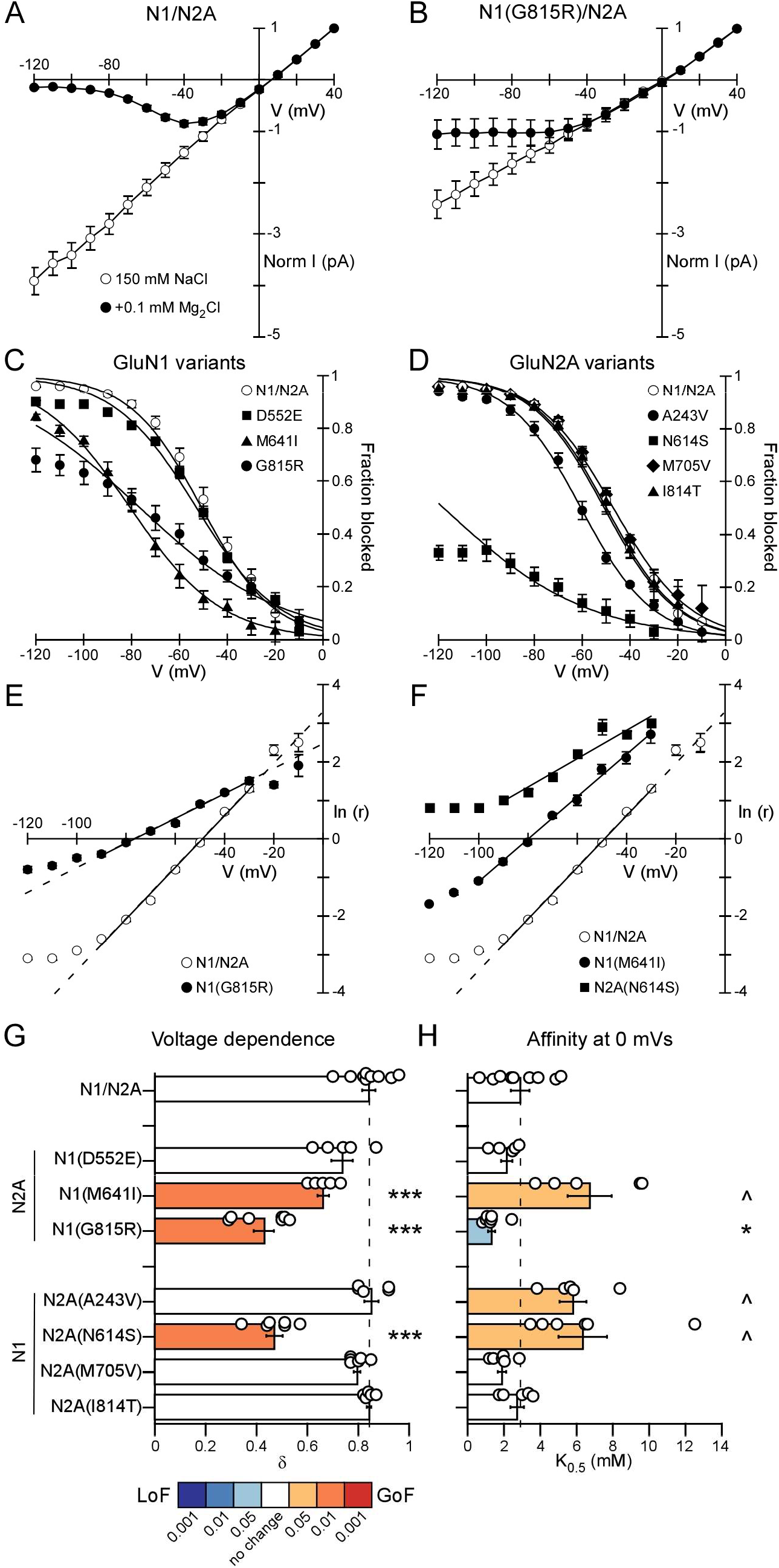
Variants alter magnesium block. (**A & B**) Peak current amplitudes (mean ± SEM) for N1/N2A (**A**) or N1(G815R)/N2A (**B**) recorded either in a 150 mM NaCl-based solution without added Mg^2+^ (I_0_, open circles) or in the same solution with added 0.1 mM MgCl2 (I_B_, filled circles). Currents were elicited by a 200 ms application of glutamate (1 mM). Peak current amplitudes were normalized to those at +40 mV. (**C & D**) Fraction blocked (mean ± SEM) for wild-type N1/N2A (open circles) and the GluN1 (**C**) or GluN2A (**D**) variants. Continuous curves are fits of Equation 1 using parameters from Figures 4G & 4H and Table 5. (**E & F**) Quantifying the voltage dependence (δ) and affinity at 0 mVs (K_0.5_(0 mV)) of Mg^2+^ block by linearizing I_B_ and I_0_ (Eqn. 2). Mean (± SEM) ln(I_p_/I_0_ – I_B_)), referred to as ln(r), plotted against voltage for 0.1 mM Mg^2+^ for wild-type (open circles) and indicated variants (solid symbols). Fits were made over the apparent linear region from −90 to −30 mV (wild-type, N1(G815R), & N2A(N614S)) and −100 to −50 mV (N1(M641I)). The analysis was restricted to ln(r) values less than 2.5 where the current amplitude was reduced by at least 10%. (**G & H**) Bar graphs (mean ± SEM) showing δ (**G**) or K_0.5_(0 mV) (**H**). Values are significantly less (*p < 0.05, ***p < 0.001) or greater (^p < 0.05) than wild-type, Student t-test (Table 5).

**Table 5.**
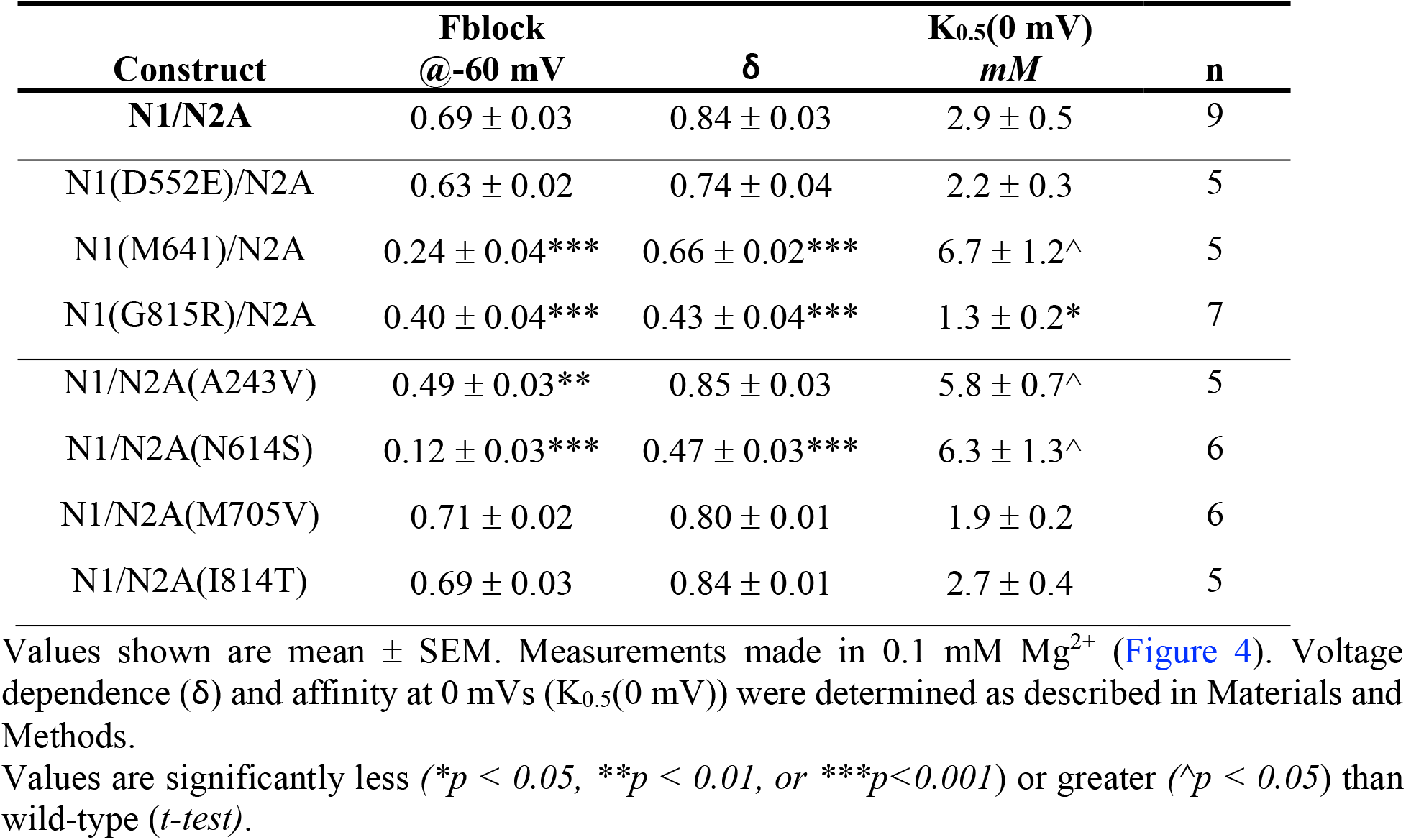
Mg^2+^ block in wild-type N1/N2A and NMDAR variants.

To assay Mg^2+^ block, we measured current amplitudes over a wide voltage range either in a solution without added Mg^2+^ (I_0_) or the same solution with added 0.1 mM Mg^2+^ (I_B_) (Figures 4A & 4B). We used this moderate concentration of Mg^2+^ since it permitted a more dynamic assessment of the voltage-dependence of the block (Wollmuth *et al*., 1998).

In terms of the fraction blocked, several of the variants produced a strong reduction in the block, including N1(M641I), N1(G815R) (Figure 4C), and N2A(N614S) (Figure 4D). To quantify the block, we linearized the relationship between I_B_ and I_0_ (Figures 4E & 4F) (see Material and Methods). This approach is advantageous since it allows for the quantification of δ, the voltage dependence of the block, and K_0.5_(0 mV), the affinity at 0 mVs, with a single Mg^2+^ concentration. Defining δ and K_0.5_(0 mV) allows us to calculate the block with any Mg^2+^ concentration and at any voltage.

With this linearization approach, the linear region of ln(r), where r = I_B_/(I_0_ – I_B_), follows the Woodhull model. For wild-type, this linear region typically extended from −90 to −30 mVs (Figure 4E, solid line fit to open circles) (Wollmuth *et al*., 1998). The deviation at more negative potentials (<−90 mV), which would be outside of a typical physiological range, may reflect a weak Mg^2+^ permeation. At more positive potentials (−20 & −10 mVs), the block was too weak and variable to be fitted in individual records, but on average often followed the extrapolated fit (Figure 4E, dashed lines). For wild-type under these conditions, δ was around 0.84 (0.84 ± 0.03, n = 9) and K_0.5_(0mV) was around 2.9 mM (2.9 ± 0.5 mM) (Table 5), consistent with previous reports (Wollmuth et al., 1998).

Quantitatively, the variants that altered Mg^2+^ block typically either decreased the voltage dependence (Figure 4G) and/or attenuated the affinity at 0 mVs (Figure 4H). The exception is N1(G815R) which attenuated the voltage dependence but enhanced affinity (Figure 4C). Although we do not know the basis of these effects, they are not surprising given that the GluN1 M4 segment regulates the topology of the M2 loop (Amin et al., 2018). Disruption of the block by N1(M641I) and N2A(N614S) is not surprising given their location in the outer vestibule and the M2 pore loop, respectively (Figure 1) (Wollmuth et al., 1996; Li et al., 2019; Xu et al., 2021). Surprisingly, N2A(A243V), which is located in the extracellular ATD, did not affect the voltage dependence but did alter the affinity at 0 mVs.

### Charge transfer over a wide voltage range

Our next goal was to derive a single parameter that would account for the effect of NMDAR activation on membrane excitability. We will refer to this parameter as ‘charge transfer’. This term encompasses variations in the current integral (Table 3), which reflects differences in membrane expression and the ability to encode synaptic activity (differences in gating) and over a wide voltage range to encompass changes in Mg^2+^ block.

To derive the charge transfer, we started with the normalized Q-V relationships as a 0 mM Mg^2+^ reference (Figures 3 & 5A, *left panel*), and then generated Mg^2+^ block curves using average δ and K_0.5_(0 mV) at 1.0 mM Mg^2+^ (Figures 5A, *right panel*). We used 1 mM Mg^2+^ since it is an approximate physiological concentration (Kapaki et al 1989). We made such calculations for wildtype and each variant (Figure 5B, Table 4). While changes in membrane voltage add additional aspects to the block by Mg^2+^ (Kampa *et al*., 2004; Clarke & Johnson, 2006), the present characterization captures the basic features of this process.

**Figure 5.**
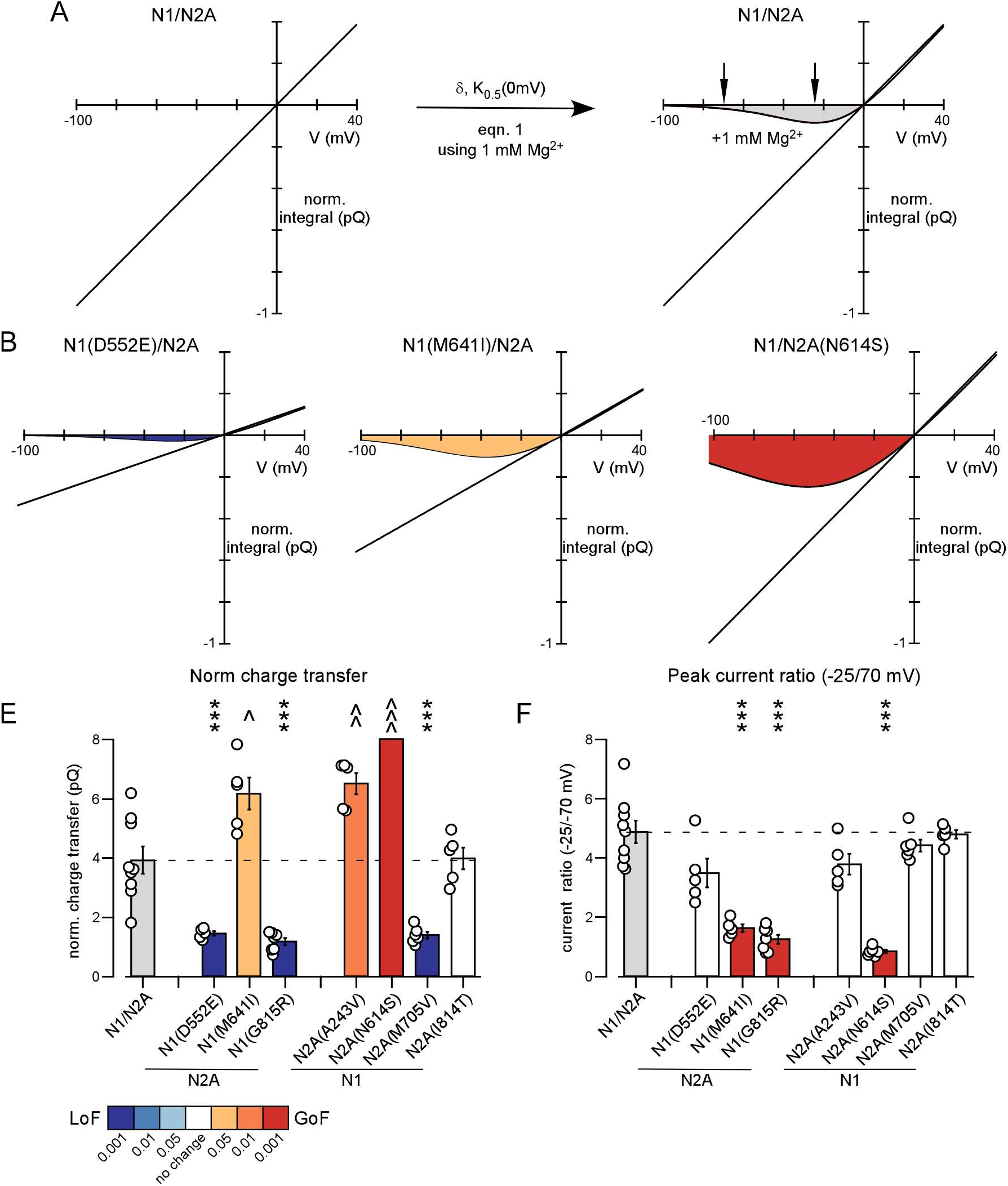
Integration of NMDAR encoding with Mg^2+^ block details charge transfer over a wide voltage range. (**A**) Integration of Mg^2+^ block with Q-V relationship for N1/N2A. *Left*, normalized Q-V in no Mg^2+^ (Q_0_) (Figure 3C). *Right*, curved line (Q_B_) is the Q-V relationship with Mg^2+^ block (1 mM). Q_B_ was derived using Eqn. 1 with Q_0_ = I_0_ and Q_B_ = I_B_ and average δ and K_0.5_(0 mV) values. Shaded area indicates charge transfer and arrows indicate −70 and −25 mVs. (**B**) Integration of Mg^2+^ block with Q-V relationship for N1(D552E), N1(M641I), and N2A(N614S). Shading reflects LoF/GoF (inset to panel E). (**E & F**). Bar graphs (mean ± SEM) showing normalized charge transfers (**E**) and ratio of current amplitudes at −25/−70 mV (**F**). Values are significantly less (***p < 0.001) or greater (^p < 0.05, ^^^ *p* < 0.001) than wild-type, Student t-test) (Table 4).

In terms of the normalized net charge transfer, wild-type was 3.9 pQ (3.9 ± 0.5 pQ, n = 9). Relative to wild-type, three variants showed a dramatic LoF: N1(D552E), N1(G815R), and N2A(M705V) (Figure 5E). Since N1(D552E) and N2A(M705V) did not significantly alter Mg^2+^ block, this LoF reflects their significant decrease in the physiological integrals (Figure 2C). N1(G815R) did show an altered Mg^2+^ block albeit this was mixed in terms of LoF and GoF, but the net effect, given the strong reduction in physiological integrals, was a reduced net charge transfer.

Three constructs showed a GoF in the total charge transfer: N1(M641I), N2A(A243V), and N2A(N614S) (Figure 5E). N1(M641I) has a significantly reduced physiological integral (Figure 2C), but the strongly reduced Mg^2+^ block led to a net increased charge transfer. N2A(A243V) and N2A(N614S) had no significant change in physiological charge transfer, therefore the increase in net charge transfer reflects a weak (N2A(A243V)) or a strong (N2A(N614S)) reduction in Mg^2+^ block.

To further understand how these variants might affect NMDAR-mediated signaling, we also characterized the ratio between the current amplitude at −25 mVs, where the inward transfer is near a peak for wild-type, and −70 mV, a typical resting membrane potential (arrows in Figure 5A, *right panel*). Current flow at −70 mV would be an index of the ‘leakage current’ at rest. Wild-type showed a −25/−70 mV ratio of 4.9 (4.9 ± 0.4, n = 9). N1(M641I), N1(G815R) and N2A(N614S) showed a significant decrease in this ratio (Figure 5F), reflecting predominantly an increased leakage current at the putative resting potential.

In summary, subsets of the variants significantly alter net charge transfer, that is the influx of both Na^+^ and Ca^2+^, in the presence of physiological Mg^2+^. Often this reflects a significant change in membrane expression (Figure 2C), but in one case, GluN1(M641I), charge transfer is flipped from LoF to GoF due to a significant reduction of Mg^2+^ block.

### NMDAR variants alter Ca^2+^ influx

As a next step, we determined the effect of the variants on Ca^2+^ influx using fractional Ca^2+^ currents (P_f_s) (Neher, 1995) (see Materials and Methods). P_f_s are advantageous over other approaches to quantify Ca^2+^ influx (e.g., changes in reversal potentials), since they can be measured under physiological conditions, can be assayed over a wide voltage range, and do not require Goldman-Hodgkin-Katz (GHK) assumptions (Jatzke et al., 2002).

To quantify P_f_s, we recorded currents and associated changes in F_385_ in 1.8 mM Ca^2+^ at −60 mVs compared to the reversal potential (Figure 6A, Table 6), though for N1(G815R) we used 5 mM Ca^2+^ because of its strong effect on Ca^2+^ influx and low amplitude (Amin et al., 2018). Under these conditions, wild-type N1/N2A showed a P_f_ of around 16% (16.2 ± 1.0%, n = 11). Two variants, N1(D552E) and N2A(I814T), showed a significant increase in Ca^2+^ influx (Figure 6B), whereas two, N1(M641I) and N1(G815R), showed significant decreases (Figures 6B & 6C). That N1(M641I) and N1(G815R) significantly affect Ca^2+^ influx is not surprising given that M641I is located in the vestibule outside of the M2 pore loop and G815R has been shown previously to affect Ca^2+^ permeability (Amin et al., 2018). On the other hand, N1(D552E) and N2A(I814T) enhancing Ca^2+^ influx is somewhat surprising. However, these positions are located in the short LBD-TMD linker regions and may in some way affect the access of Ca^2+^ for the pore. In any case, these results highlight variants outside of the M2 pore loop can impact Ca^2+^ influx.

**Figure 6.**
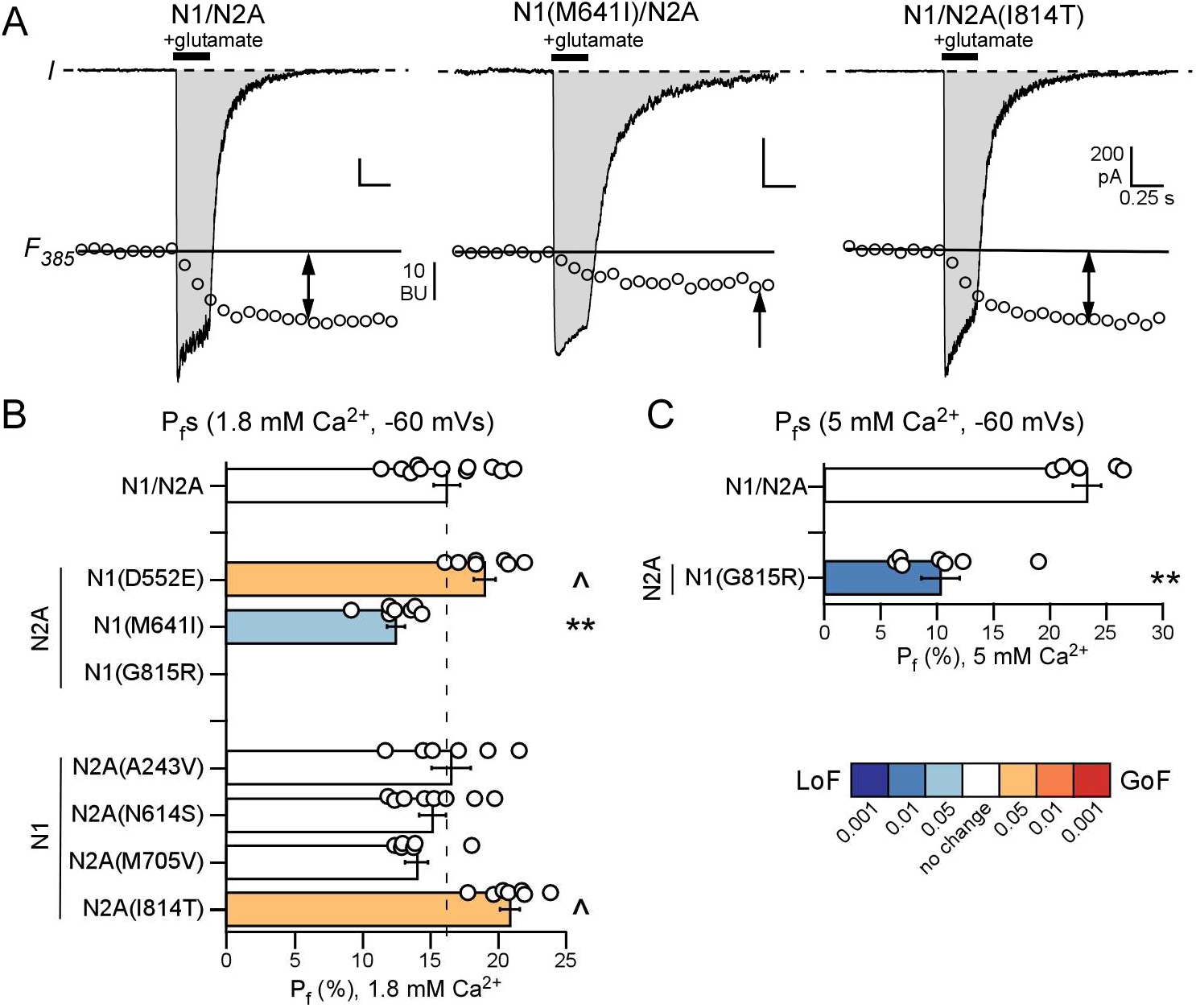
Quantification of fractional Ca^2+^ currents (P_f_s). (**A**) Simultaneous measurements of whole-cell currents (*I*) and fluorescence intensity with 385 nm excitation (*F_385_*) evoked by glutamate (filled bars) in HEK293 cells expressing N1/N2A (*left panel*), N1(M641I) (*middle panel*), or N2A(I814T) (*right panel*). The potential (V) was −60 mV to the reversal potential (see Methods). In the current records, the shaded regions correspond to the current integral (Q_T_). F_385_ is expressed in bead units (BU). ΔF_385_ was derived as the difference between the F_385_ amplitude at the indicated time (arrows) and the baseline F_385_ signal (continuous line), extrapolated from a linear fit to the F_385_ amplitudes prior to the glutamate application. Q_T_ is approximately the same for each of the example records. (**B & C**) Bar graph (mean ± SEM) showing average P_f_ for N1/N2A or variants at −60 mVs either in 1.8 mM (**B**) or 5 mM (**C**) Ca^2+^. Values are significantly less (*\*\*p < 0.01*) or greater (*^p < 0.05*) than wild-type, *Student t-test* (Table 6).

**Table 6.** Fractional Ca^2+^ current in wild-type N1/N2A and NMDAR variants.

| Extra.<br>$\text{Ca}^{2+}$ | Construct | $P_f$<br>%<br>-60 mV | $P_f$<br>%<br>-20 mV | Ratio<br>$P_{f,-20}/P_{f,-60}$ | $P_{\text{Ca}}/P_{\text{Na}}$ | n |
| --- | --- | --- | --- | --- | --- | --- |
| **1.8 mM** | **N1/N2A** | $16.2 \pm 1.0$ | $20.2 \pm 1.1$ | $1.27 \pm 0.02$ | $3.8 \pm 0.3$ | 11 |
| | N1(D552E)/N2A | $19.0 \pm 0.8^{\wedge}$ | $23.9 \pm 1.1^{\wedge}$ | $1.25 \pm 0.03$ | $4.5 \pm 0.2^{\wedge}$ | 7 |
| | N1(M641I)/N2A | $12.5 \pm 0.7^{**}$ | $14.9 \pm 0.7^{**}$ | $1.20 \pm 0.02^*$ | $2.8 \pm 0.2$ | 7 |
|  | N1(G815R)/N2A | -- |  |  |  |  |
| | N1/N2A(A243V) | $16.6 \pm 1.4$ | $21.5 \pm 1.7$ | $1.30 \pm 0.03$ | $3.9 \pm 0.4$ | 6 |
| | N1/N2A(N614S) | $15.2 \pm 1.0$ | $19.1 \pm 1.1$ | $1.29 \pm 0.04$ | $3.5 \pm 0.3$ | 8 |
| | N1/N2A(M705V) | $14.0 \pm 0.9$ | $18.2 \pm 1.5$ | $1.28 \pm 0.03$ | $3.2 \pm 0.2$ | 6 |
| | N1/N2A(I814T) | $20.9 \pm 0.7^{\wedge\wedge}$ | $27.6 \pm 1.0^{\wedge\wedge\wedge}$ | $1.33 \pm 0.04$ | $5.1 \pm 0.2$ | 7 |
| **5 mM** | **N1/N2A** | $23.3 \pm 1.2$ | $37.1 \pm 2.8$ | $1.60 \pm 0.11$ | $2.1 \pm 0.2$ | 5 |
| | N1(G815R)/N2A | $10.3 \pm 1.7^{***}$ | $15.2 \pm 2.0^{***}$ | $1.52 \pm 0.06$ | $0.8 \pm 0.15^{***}$ | 7 |

Values shown are mean ± SEM. $P_f$ for G815R was determined in 5 mM $\text{Ca}^{2+}$ due to low current amplitudes and low $\text{Ca}^{2+}$ permeability (Amin *et al.*, 2018).
$P_{\text{Ca}}/P_{\text{Na}}$ was calculated using eq 5 (see Materials and Methods).
Values are significantly less ( $**p < 0.01$ or $***p < 0.001$ ) or greater ( $^{\wedge}p < 0.05$ , $^{\wedge\wedge}p < 0.01$ , or $^{\wedge\wedge\wedge}p < 0.001$ ) than the respective wild-type (*t-test*).

### Integrating Ca^2+^ influx over a wide voltage range

To generate a more dynamic view of Ca^2+^ influx in wild-type and the variants, we wanted to define it during trains of glutamate pulses and over a wide voltage range. To do so, required two key considerations. First, we needed to verify that fractional Ca^2+^ currents were the same with long glutamate applications and with trains of glutamate pulses. Nevertheless, these experiments were challenging since the fast glutamate application system with trains of pulses compromised the optics of measuring Ca^2+^ influx. Nevertheless, we were able to record in a limited number of instances and within the same cell fractional Ca^2+^ currents with long pulses and trains of glutamate applications (e.g., Figures 7A & 7B). For both wild-type and the variants, we could detect no difference between long applications and glutamate pulses (Table 7). Hence, we can use the fractional Ca^2+^ currents measured using long pulses to characterize Ca^2+^ influx with trains of glutamate pulses (Figure 7C).

**Figure 7.**
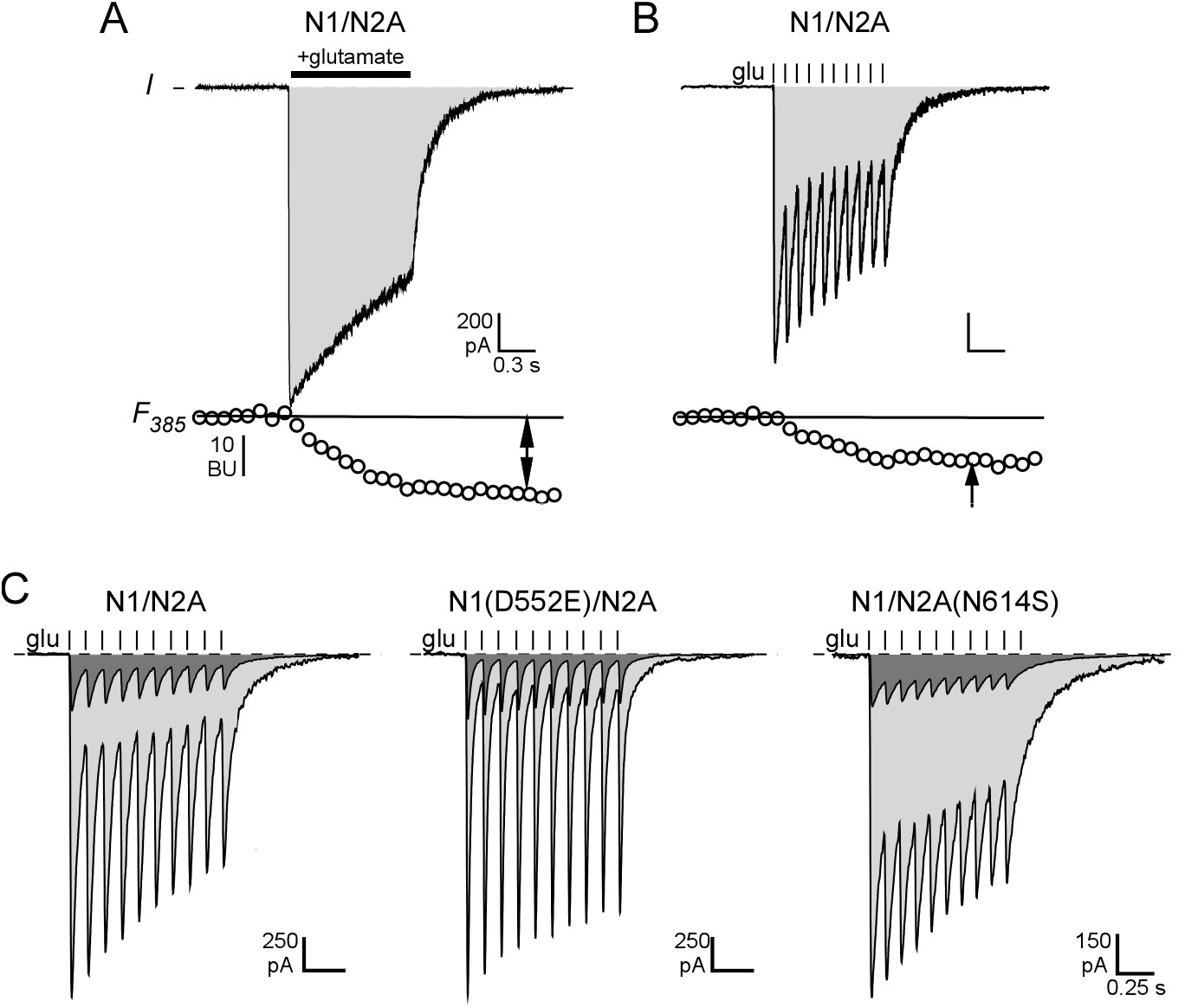
Ca^2+^ influx during trains of glutamate pulses. (**A & B**) Fractional Ca^2+^ currents are the same whether evoked by continuous or pulsatile glutamate pulses. Simultaneous measurements of whole-cell currents (I, top) and fluorescence intensity with 385 nm excitation (*F_385_*, bottom) evoked by either continuous glutamate (1000 s) (**A**) or 10 glutamate pulses at 10 Hz (**B**) (Table 7). Recordings were made sequentially within the same cell, but only recordings where the baseline *F_385_* was within 10% were included in analysis. (**C**) Pulses of glutamate (from Figure 2A, light gray), showing a representation of the fraction of charge carried by Ca^2+^ (dark gray). Original pulse is reduced by average P_f_ at −60 mV (Figure 6).

**Table 7.** Calcium charge transfer during continuous and pulsatile currents in wild-type N1/N2A and NMDAR variants.

| Construct | Ratio Continuous/<br>Pulse | n | Normalized Ca<sup>2+</sup><br>charge transfer<br>(pQ) | n |
| --- | --- | --- | --- | --- |
| N1/N2A | 0.99 ± 0.03 | 5 | 0.78 ± 0.04 | 11 |
| N1(D552E)/N2A | 1.00 ± 0.03 | 3 | 0.32 ± 0.01*** | 7 |
| N1(M641I)/N2A | 0.97 ± 0.02 | 4 | 0.83 ± 0.04 | 7 |
| N1(G815R)/N2A | Not tested |  | 0.08 ± 0.01*** | 7 |
| N1/N2A(A243V) | 0.88 ± 0.04 | 3 | 1.0 ± 0.1 | 6 |
| N1/N2A(N614S) | 1.06 ± 0.05 | 3 | 1.8 ± 0.1^^^ | 8 |
| N1/N2A(M705V) | 0.99 ± 0.09 | 3 | 0.23 ± 0.01*** | 6 |
| N1/N2A(I814T) | 0.93 ± 0.04 | 4 | 0.72 ± 0.02 | 7 |

The second issue is to characterize Ca^2+^ influx over a wide voltage range, not just at −60 mVs. While P_f_s could be measured over a wide voltage range for wild-type and each variant, such experiments would be extremely time-consuming. We therefore took advantage of previous observations that P_f_s follow GHK assumptions at physiological concentrations of Ca^2+^ (Schneggenburger, 1996; Jatzke et al., 2002), allowing P_Ca_/P_Na_ to be calculated which in turns allows P_f_s to be calculated over a wide voltage range (Eqn. 5, Materials & Methods). To validate this idea, we measured P_f_s over a wide voltage range for wild-type (Figure 8A), and verified that human N1/N2A could be characterized by a single P_Ca_/P_Na_ (solid line), which was 3.8 (3.8 ± 0.3, n = 11). For each variant, we measured P_f_s at −60 and −20 mVs and used the ratio of these values to verify that they follow GHK assumptions (Table 6). All variants except for N1(M641I) showed no significant difference from wild-type. A similar observation was made for N1(G815R) using 5 mM Ca^2+^, as shown previously P_Ca_/P_Na_s deviate from GHK at concentrations away from 1.8 mM (Jatzke et al., 2002).

**Figure 8.**
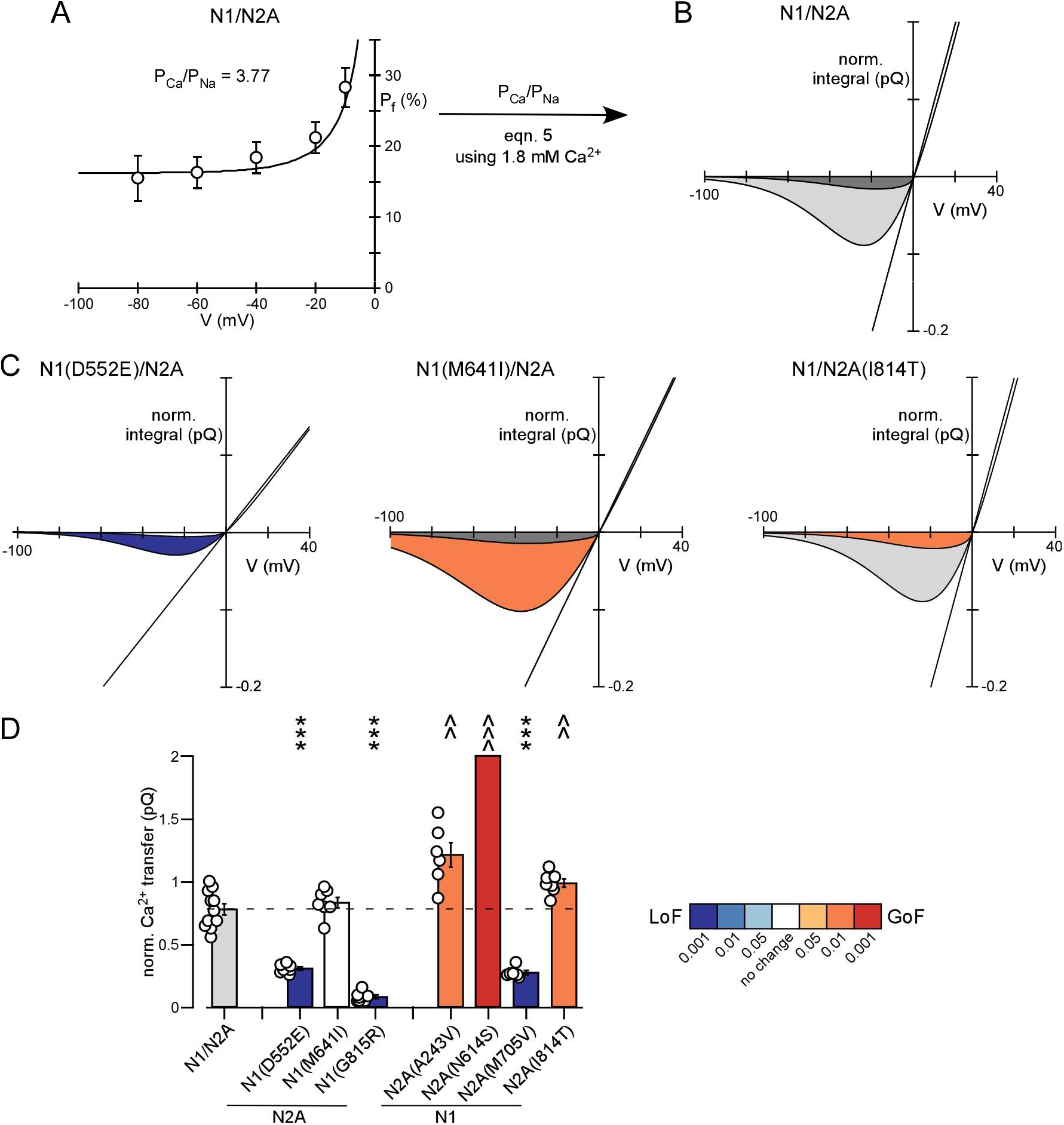
Integration of net charge transfer and fractional Ca^2+^ currents define Ca^2+^ influx over a wide voltage range. (**A**) Human GluN1/GluN2A follow GHK at 1.8 mM Ca^2+^. Voltage dependence of P_f_ values (mean ± SEM). Continuous line is predicted P_f_ values for a single P_Ca_ value, using Eqn. 5, with P_Ca_/P_Na_ derived from the P_f_ measurements at −60 mV (Table 6). (**B**) Integration of charge transfer (light gray from Figure 5A) and Ca^2+^ influx (dark gray) for N1/N2A using the fractional Ca^2+^ current from the derived P_Ca_/P_Na_ (= 3.8 for wild-type). (**C**) Average charge and Ca^2+^ transfer for N1(D552E), N1(M641I), and N2A(I814T). Integrals are shown as a shade matching LoF/GoF or a shade of gray if not different from wild type. For N1(D552E), both the charge and Ca^2+^ (shaded white) were significantly less than wild-type. (**D**) Bar graph (mean ± SEM) showing normalized total Ca^2+^ transfer (**D**). Values are significantly less (*\*p<0.05, **p<0.01, ***p < 0.001*) or greater (*^^^p < 0.001*) than wild-type, *Student t-test* (Table 7).

### Effect of variants on Ca^2+^ influx over a wide voltage range

To compare Ca^2+^ influx over a wide voltage range between wild-type and the variants, we generated a single parameter, termed ‘Ca^2+^ transfer’. To derive this term, we used P_Ca_/P_Na_ estimated from the voltage dependence of P_f_ (Figure 8A; Table 6) and Eq. 5 to calculate P_f_s over a continuous voltage range. Then using the currents in the presence of Mg^2+^ as a reference (Figure 8B, *light shade*), we used these P_f_s, again which are a percentage, to derive the percent of the total current carried by Ca^2+^ over a wide voltage range (Figure 8B, *dark shade*). Ca^2+^ transfer is the integral (Figure 8B, *dark shade*) of this current from −100 to 0 mVs. We made similar calculations for all variants (Figure 8C).

Three variants, N1(D552E), N1(G815R), and N2A(M705V), showed significant decreases in net Ca^2+^transfer (Figure 8D). Notably, N1(M641I) showed a GoF in terms of charge transfer, but its net Ca^2+^ transfer was indistinguishable from wild-type (Figure 8D), reflecting that it showed a LoF in terms of fractional Ca^2+^ currents (Figure 6B). Variants N2A(A243V), N2A(N614S) and N2A(I814T) showed to varying degrees a GoF in terms of Ca^2+^ transfer.

### Negative allosteric modulation to normalize charge and Ca^2+^ transfer in GoF variants

Clinically, the goal is to ‘normalize’ LoF and GoF variants back to wild-type to recover as much function as possible. To normalize net charge and Ca^2+^ transfer for two GoF variants, N1(M641I) and N2A(N614S), we treated wild-type and these two variants with Dextrorphan (DEX), a negative allosteric modulator (NAM) of NMDARs (Pierson *et al*., 2014) (Figure 9). To assay the effects of DEX, we used our standard pulse protocol (Figure 9A). Wild-type and both variants showed a decrease in peak amplitude (Figure 9B) and charge transfer (Figures 9C & 9D). As found previously (Xu *et al*., 2021), DEX is more efficacious at N1(M641I), so we assayed two concentrations, 3 μM (Figures 9B & 9C) and 0.3 μM (Figure 9D).

**Figure 9.**
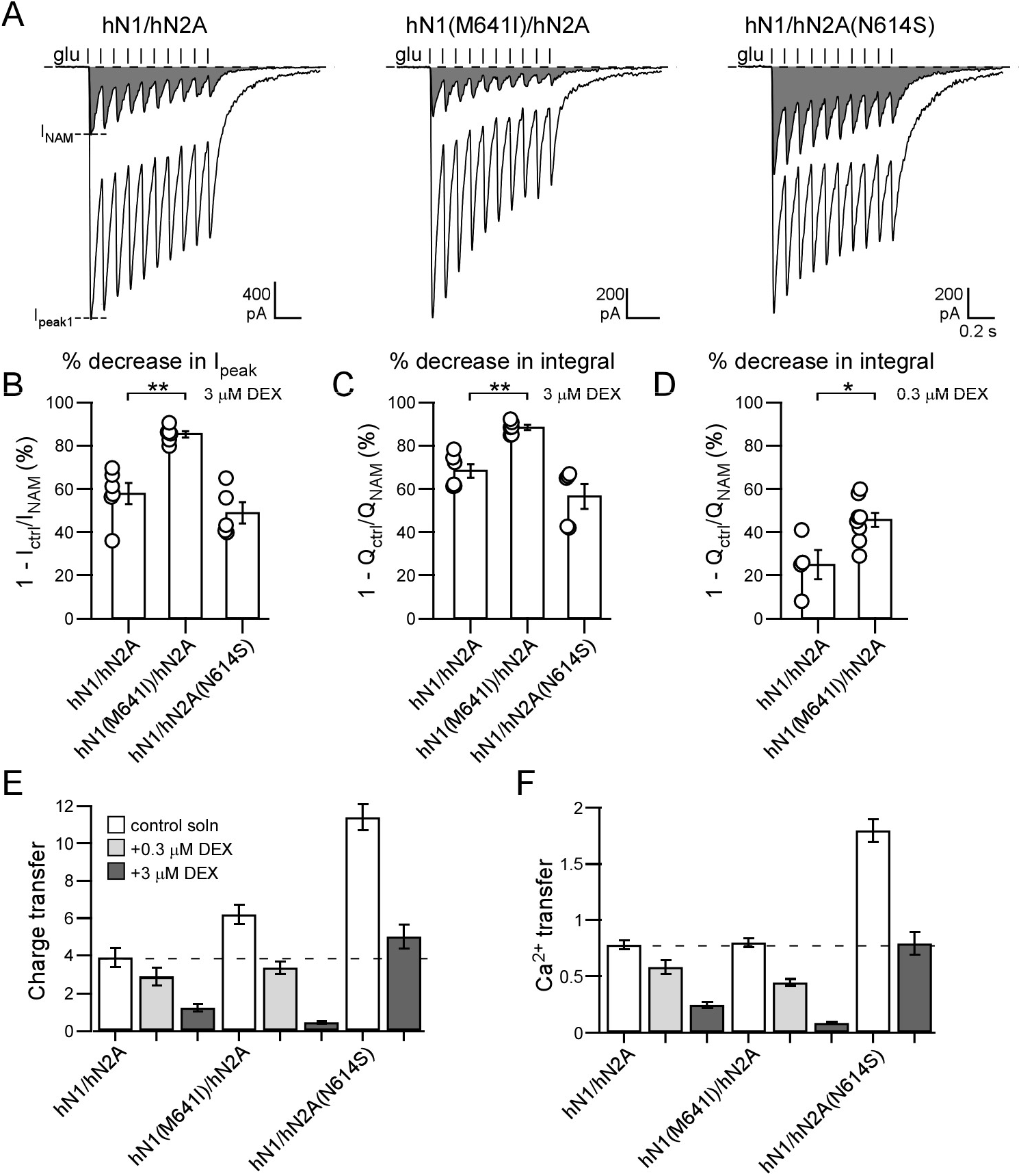
A negative allosteric modulator normalizes charge and Ca^2+^ transfer in GoF variants to wild-type levels. (**A**) Whole-cell currents elicited by 10 glutamate pulses at 10 Hz in the absence (I_peak1_, white) or presence (I_NAM_, gray) of 3 μM dextrorphan (DEX), a negative allosteric modulator (NAM) of NMDARs. Recordings are displayed and analyzed as in Figure 2A. (**B-D**) Bar graphs (mean ± SEM) showing % decrease in I_peak1_ (**B**) or charge transfer normalized to membrane capacitance without (Q_ctrl_) and with (Q_NAM_) DEX for 3 μM DEX (**C**) or 0.3 μM DEX (**D**). For 3 μM DEX, the number of recordings, from left to right: 6, 6, 5; For 0.3 μM DEX, from left to right: 4, 8. (**E & F**) Bar graphs showing charge transfer (**E**) or Ca^2+^ transfer **(F)** in control solution or in DEX. See inset for conditions. Values were calculated based on the % decrease in current integral. Values are significantly less (*\*p < 0.05, **p < 0.01*) than wild-type, *Student t-test*.

Application of 3 μM DEX reduced N2A(N614S) charge (Figure 9E) and Ca^2+^ (Figure 9F) transfers to near wild-type values. On the other hand, results with N1(M641I) were more complex since while charge transfer shows GoF (Figure 9E), Ca^2+^ transfer does not (Figure 9F). In addition, N1(M641I) showed an increased sensitivity to DEX (Figures 9B-9D) (Xu *et al*., 2021). Accordingly, 3 μM DEX nearly eliminated any response from N1(M641I), whereas 0.3 μM DEX normalized charge transfer back to wild type (Figure 9E) but reduced Ca^2+^ transfer below wild type levels (Figure 9F).

### Positive allosteric modulation to normalize charge and Ca^2+^ transfer in LoF variants

To normalize net charge and Ca^2+^ transfer for two LoF variants, N1(D552E) and N2A(M705V), we treated wild-type and these two variants with GNE-9278, a non-specific positive allosteric modulator (PAM) of NMDARs (Wang et al., 2017) (Figure 10). Notably, GNE-9278 enhanced peak current amplitudes for all constructs to about the same levels (Figures 10A & 10B). However, it also significantly slowed deactivation rates in wild-type (210 ± 10 ms to 2430 ± 850 ms, n = 4) and in N2A(M705V) (240 ± 20 ms to 2240 ± 690 ms, n = 5), but not in N1(D552E) (100 ± 15 ms to 120 ± 18 ms, n = 5). Because of this slowing, peak amplitudes were basically lost during the pulse train (Figure 10A), and the current integral was more strongly potentiated (Figure 10C) than peak current amplitudes. Correspondingly, charger transfer was nearly 2 times larger than changes in peak amplitude for wild-type and N2A(M705V), whereas in N1(D552E) it was about the same (Figure 10D).

**Figure 10.**
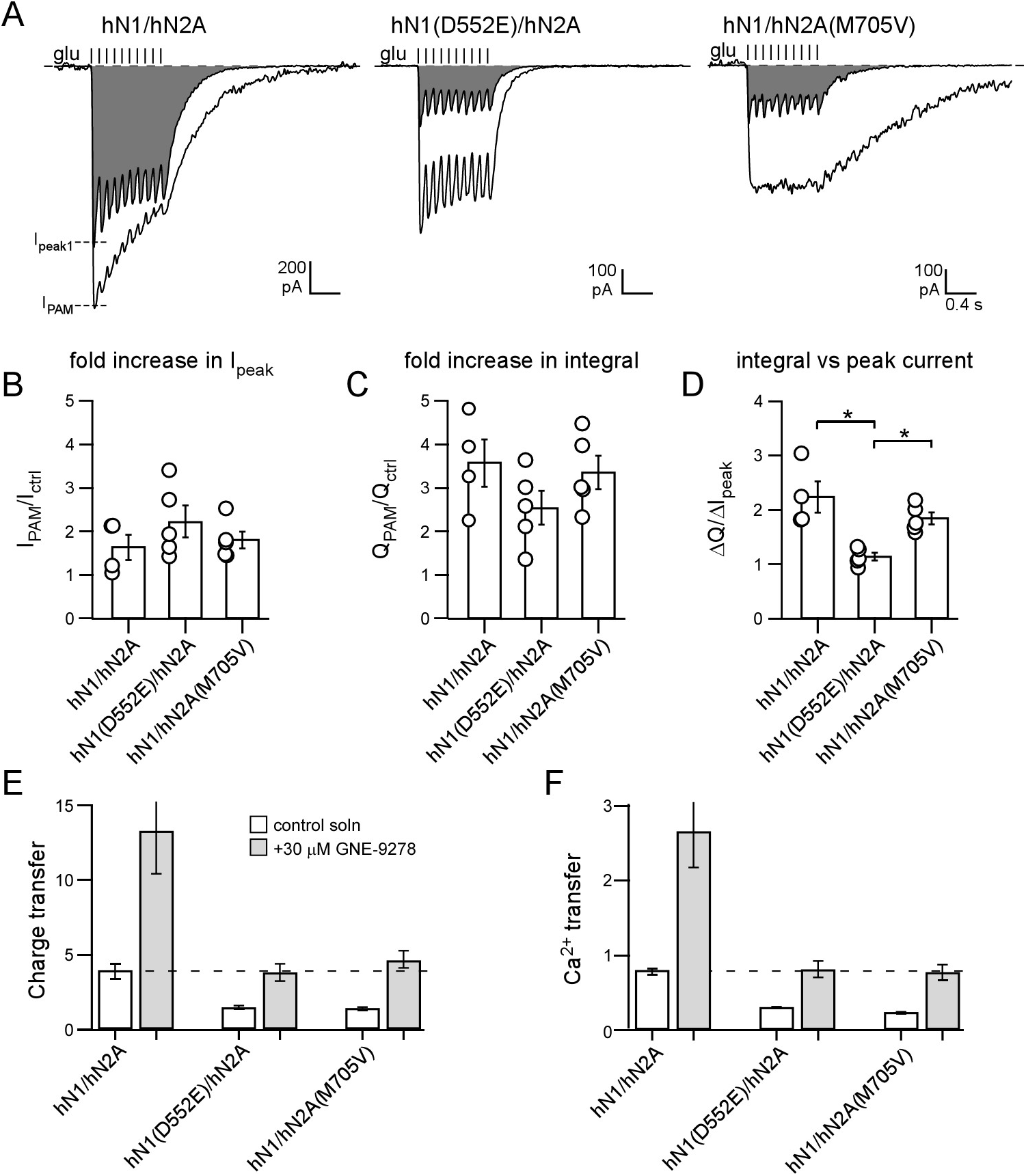
A positive allosteric modulator normalize charge and Ca^2+^ transfer in LoF variants to wild-type levels. (**A**) Whole-cell currents elicited by 10 glutamate pulses at 10 Hz in the absence (I_peak1_, gray) or presence (I_PAM_, white) of 30 μM GNE-9278, a positive allosteric modulator (PAM) of NMDARs. Recordings are displayed and analyzed as in Figure 2A though we show an extended time base so that the GNE-9278-induced slowing of deactivation in wild-type and M705V could be seen. (**B-C**) Bar graphs (mean ± SEM) showing fold-change in I_peak1_ (**B**) or charge transfer (**C**) in the presence of GNE-9278 relative to its absence. The number of recordings, from left to right: 4, 5, 5. None of the values are significantly different. (**D**) Ratio of the GNE-9278-induced change in charge transfer relative to peak current amplitude. *\*p < 0.05, one-way ANOVA with post-hoc Tukey’s test*. (**E & F**) Bar graphs showing charge transfer (**E**) or Ca^2+^ transfer **(F)** in control solution or in GNE-9278. See inset for conditions. Values were calculated based on the fold increase in charge transfer.

For both variants, 30 mM GNE-9278 was able to normalize charge (Figure 10E) and Ca^2+^ (Figure 10F) transfer back to wild type levels.

## DISCUSSION

At synapses, NMDARs use a variety of biophysical and molecular mechanisms to encode presynaptic activity and regulate their signaling (Hansen et al., 2021). To characterize NMDAR disease-associated variants, we devised an approach that encompasses many aspects of NMDAR function – membrane expression, diverse gating properties, Mg^2+^ block, and Ca^2+^ influx – into two parameters: ‘charge transfer’ (Figure 5E) and ‘Ca^2+^ transfer’ (Figure 8D). These parameters were integrated over a wide physiological voltage range. Charge transfer encompasses how NMDARs will affect membrane excitability, whereas Ca^2+^ transfer encompasses the magnitude of Ca^2+^ influx. These parameters do not account for the full properties of NMDARs at synapses (see below) but do encompass many of them and allow for high-throughput analysis of patient variants. Importantly, these parameters represent a broader target to normalize using either NAMs for GoF variants (Figure 9) or PAMs for LoF variants (Figure 10).

### Integrated approach to assay NMDAR function

The role of ion channels within a circuit is complex and difficult to explain in a heterologous system, especially for receptors with complex regulation and functions such as the NMDAR. The impact of NMDARs on brain function depends on three general considerations: (i) a charge transfer and Ca^2+^-mediated signaling that arises from the glutamate-induced opening of the associated ion channel (Paoletti et al., 2013; Wollmuth, 2018; Hansen et al., 2021); (ii) a metabotropic pathway that signals independently of ion channel opening (Nabavi et al., 2013; Valbuena & Lerma, 2016; Rajani et al., 2020); and (iii) NMDAR cell biology, which encompasses subunit composition and post-translational modifications as well as the number and distribution of NMDARs on the membrane (Paoletti et al., 2013; Lussier et al., 2015; Groc & Choquet, 2020).

Neurons in the brain rarely convey information via a single action potential but rather show a wide range of activity from several Hz to 100s of Hz (Buzsaki & Mizuseki, 2014). The synaptic output of this firing will depend on the release probability of presynaptic terminals. As a reference, we used 10 pulses at 10 Hz (e.g., Figure 2 & Table 3), which would approximate a neuron firing at 10 Hz with a release probability of 1 to one firing at 100 Hz with a release probability of 0.1. While many different numbers and frequencies could be tested, this pulse protocol captures the dynamics of NMDARs at synapses.

A notable feature of using a pulse protocol is that it integrates NMDAR gating including rates of activation, deactivation, and desensitization and recovery from desensitization into a single ‘integral’ term (Figure 2 & Table 3). Highlighting this integration of gating properties are the outcomes for GluN1(D552E), GluN1(M641I), and GluN2A(N614S). In general, one would anticipate that the rate of deactivation would be proportional to the current integral. Indeed, relative to wild-type, GluN1(D552E) shows a faster deactivation rate and correspondingly a reduced integral, whereas GluN2A(N614S) shows a slower deactivation rate and correspondingly a greater integral (Figure 2 & Table 3). On the other hand, GluN1(M641I) shows a slower deactivation rate, yet shows a reduced integral. This reduced integral with slower deactivation presumably reflects the enhanced desensitization displayed by this variant (Table 3). While we did not explore further the mechanistic basis for variations in integrals with rapid pulses, this outcome highlights how this pulse protocol captures diverse gating properties of NMDARs into a single term.

Using the 10 pulses at 10 Hz as a reference, we integrated the current integral over a wide voltage range with Mg^2+^ block to derive ‘charge transfer’ (Figures 3-5 & Tables 4 & 5) and then integrated these outcomes with Ca^2+^ influx to derive ‘Ca^2+^ transfer’ (Figures 6-8 & Tables 6 & 7). This approach is similar but expands on that developed by others (Swanger *et al*., 2016).

Notably our approach extends over a wide voltage range and encompasses Ca^2+^ influx. While NMDAR-mediated Ca^2+^ influx is a key functional property, few papers address it in terms of variants (Li et al., 2019). Nevertheless, unexpected variants outside the direct permeation pathway formed by M3 and the M2 loop impact Ca^2+^ permeation including the M4 segment, GluN1(G815R), and the LBD-TMD linker region, GluN1(D552E) and GluN2A(I814T) (Figure 6 & Table 6).

Our approach is straight forward requiring only the measurement of 3 parameters: current integrals, the voltage-dependence of Mg^2+^ block, and Ca^2+^ influx. Perhaps one challenge might be the measurement of fractional Ca^2+^ currents, which is not a common technique. Although less direct, as an alternative one could use relative changes in Ca^2+^ permeability assayed using reversal potentials to assay Ca^2+^ influx to derive Ca^2+^ charge transfer.

Our snapshot of NMDARs encompasses many aspects of NMDAR function – membrane expression, gating properties, Mg^2+^ block, and Ca^2+^ influx – to derive charge transfer and Ca^2+^ influx. This protocol, however, can be readily expanded to include glutamate and glycine affinity, extrasynaptic receptors, other subunits, and modulators such as Zn^2+^ and H^+^.

### Relation of outcomes to disease phenotype

We focused on NMDAR variants that are associated with an epilepsy phenotype of varying degrees of severity (Table 1). In terms of relating outcomes for heterologously expressed NMDARs with a clinical phenotype, epilepsy is optimal because symptoms are functionally quantifiable (e.g., daily seizures reduced to weekly seizures) and this disorder harbors less severe phenotypes (Table 1). Nevertheless, relating outcomes from HEK293 to the epilepsy phenotype is extremely challenging in part because these variants typically have comorbidities including autism, schizophrenia, and intellectual and developmental delays (Burnashev & Szepetowski, 2015; Hardingham & Do, 2016; Amin et al., 2021). Still, we will try to relate the most salient features of outcomes to clinical phenotypes.

A major step in treatments and understanding how variants contribute to the clinical phenotype is determining whether variants are GoF or LoF. Nevertheless, this can depend on the specific parameter measured (e.g., GluN1(M641I)) (Table 8). In addition, GoF and LoF may not be identical across parameters. For example, in terms of charge transfer, GluN1(M641I) is a GoF whereas GluN1(G815R) is a LoF (Figure 5E & Table 8), whereas both induce a GoF in terms of the ‘leak’ current at rest (Figure 5F & Table 8), which primarily reflect a disruption in Mg^2+^ block. Interestingly, produce relatively similar phenotype with a common age of onset (Table 1), suggesting that the ‘leak’ current at rest may be a key functional component.

**Table 8.**
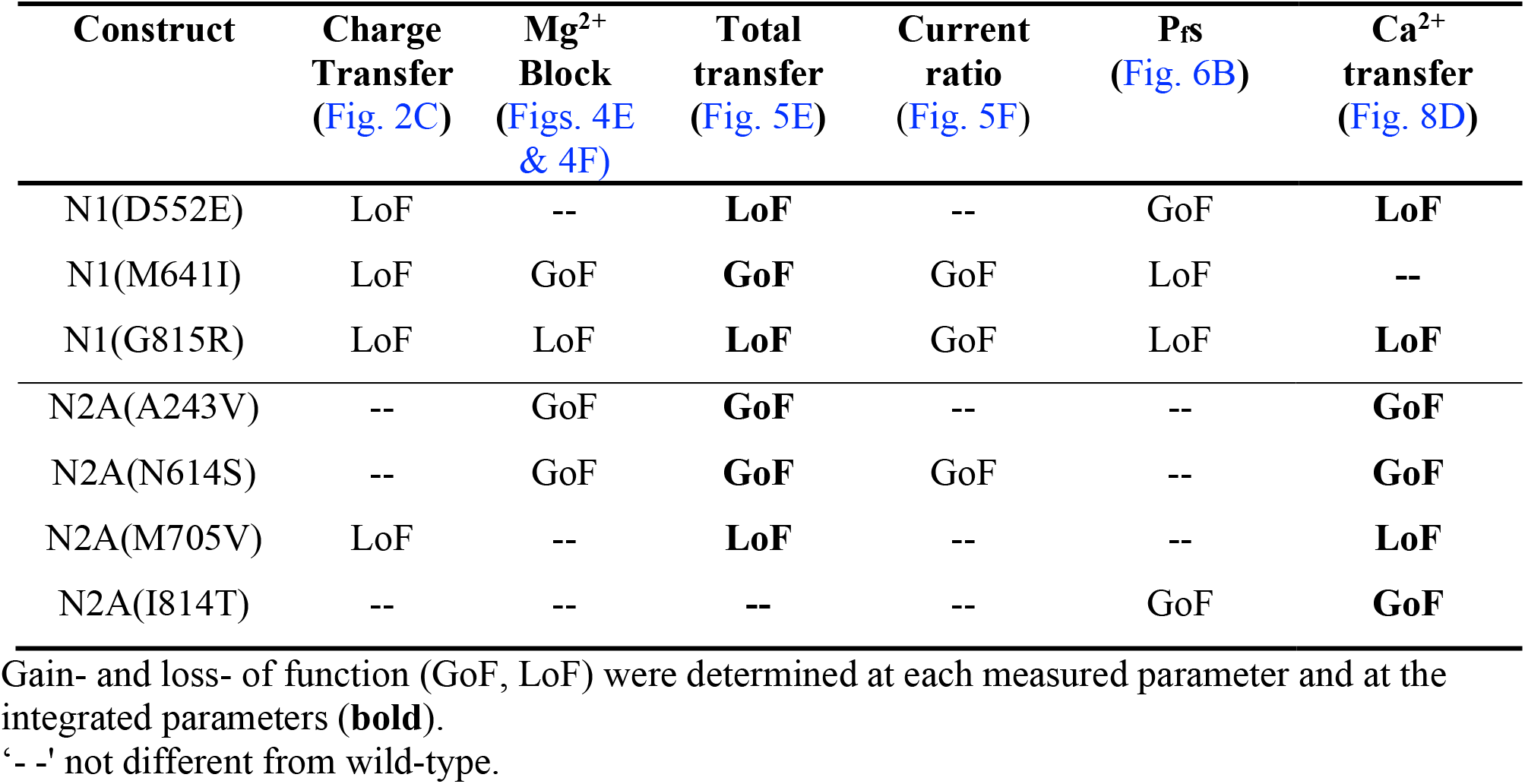
Classification of NMDAR variants as gain- or loss-of-function.

Different variants may hold clues to the consequences of misregulated calcium in terms of patient severity. I814T has no significant change in sodium handling or response to synaptic-like pulses, rather this variant shows only a change in calcium. Yet, this variant is associated with a mild epilepsy phenotype. Other assessments of this variant has completely missed this key change in calcium handling, call it a no change function. This indicates the importance of the integrated approach used here and the consequence of a small shift in NMDAR Ca handling can have symptomatically.

Variants which are more active at low potentials (i.e. −60 mV) such as M641I may result in overactive or “leaky” NMDARs which allow calcium to enter the cell in response to low-threshold signals. In epilepsy this may be a source of hyperactivity in a circuit or a driver for epileptogenesis. There is also potential for alterations in long-term potentiation or depression as calcium influx can drive one form of plasticity over the other. The ratio between sodium influx and calcium influx may also play a role in LTP versus LTD, although this is yet to be fully understood, it is altered by patient variants.

## Acknowledgments

This work was supported by a NIH RO1 from NINDS (NS088479) (LPW) and T32 GM127253 (GM).

We thank Drs. Johansen Amin and Kelvin Chan for helpful discussion, comments on the manuscript, and/or generating preliminary data for this work. Donna Schmidt is thanked for technical assistance.

## Notes

### Competing Interest Statement

The authors have declared no competing interest.

